# Intrinsic and extrinsic regulation of human fetal bone marrow haematopoiesis and perturbations in Down syndrome

**DOI:** 10.1101/2021.06.25.449771

**Authors:** Laura Jardine, Simone Webb, Issac Goh, Mariana Quiroga Londoño, Gary Reynolds, Michael Mather, Bayanne Olabi, Emily Stephenson, Rachel A. Botting, Dave Horsfall, Justin Engelbert, Daniel Maunder, Nicole Mende, Caitlin Murnane, Emma Dann, Jim McGrath, Hamish King, Iwo Kucinski, Rachel Queen, Christopher D Carey, Caroline Shrubsole, Elizabeth Poyner, Meghan Acres, Claire Jones, Thomas Ness, Rowan Coulthard, Natalina Elliott, Sorcha O’Byrne, Myriam L. R. Haltalli, John E Lawrence, Steven Lisgo, Petra Balogh, Kerstin B Meyer, Elena Prigmore, Kirsty Ambridge, Mika Sarkin Jain, Mirjana Efremova, Keir Pickard, Thomas Creasey, Jaume Bacardit, Deborah Henderson, Jonathan Coxhead, Andrew Filby, Rafiqul Hussain, David Dixon, David McDonald, Dorin-Mirel Popescu, Monika S. Kowalczyk, Bo Li, Orr Ashenberg, Marcin Tabaka, Danielle Dionne, Timothy L. Tickle, Michal Slyper, Orit Rozenblatt-Rosen, Aviv Regev, Sam Behjati, Elisa Laurenti, Nicola K. Wilson, Anindita Roy, Berthold Göttgens, Irene Roberts, Sarah A. Teichmann, Muzlifah Haniffa

## Abstract

Throughout postnatal life, haematopoiesis in the bone marrow (BM) maintains blood and immune cell production. Haematopoiesis first emerges in human BM at 12 post conception weeks while fetal liver (FL) haematopoiesis is still expanding. Yet, almost nothing is known about how fetal BM evolves to meet the highly specialised needs of the fetus and newborn infant. Here, we detail the development of fetal BM including stroma using single cell RNA-sequencing. We find that the full blood and immune cell repertoire is established in fetal BM in a short time window of 6-7 weeks early in the second trimester. Fetal BM promotes rapid and extensive diversification of myeloid cells, with granulocytes, eosinophils and dendritic cell (DC) subsets emerging for the first time. B-lymphocyte expansion occurs, in contrast with erythroid predominance in FL at the same gestational age. We identify transcriptional and functional differences that underlie tissue-specific identity and cellular diversification in fetal BM and FL. Finally, we reveal selective disruption of B-lymphocyte, erythroid and myeloid development due to cell intrinsic differentiation bias as well as extrinsic regulation through an altered microenvironment in the fetal BM from constitutional chromosome anomaly Down syndrome during this crucial developmental time window.

## Introduction

Following waves of haematopoiesis in the yolk sac (YS) and fetal liver (FL), the bone marrow (BM) is established as the site of lifelong blood and immune cell production. In humans, haematopoietic cells appear in the fetal BM from 11-12 post conception weeks (PCW)^1,2^. By this time, development of the immune repertoire has been initiated in the liver, with contributions from spleen and thymus^3,4^. The contribution of the BM, both in terms of fetal haematopoiesis and in laying foundations for BM haematopoiesis in the longer term remains to be established.

Haematopoiesis must supply the fetus with erythrocytes for oxygen transport, platelets for haemostasis, macrophages for tissue remodelling and an immune system that is poised to respond to insult without risking tissue damage. In fetal life, the priorities are to establish an innate immune system primed to respond to any pathogen encountered in perinatal life, and to provide the pool of B- and T-lymphocyte progenitors needed to prescribe the adaptive immune response to pathogens after birth, particularly in the first few weeks of life when the newborn infant is first exposed to the extra-uterine environment.

Perturbations in haematopoiesis *in utero* can have far reaching implications. Oncogenic events acquired in haematopoietic cells during fetal life, such as *ETV6-RUNX1* or *KMT2A* fusion genes, often progress to childhood leukaemia^5–8^. Somatic, *in utero GATA1* mutations leading to neonatal preleukemia and increased risk of myeloid leukaemia in early childhood are specific to Down syndrome (DS)^9,10^. Extensive abnormalities in haematopoiesis have been documented in DS FL preceding *GATA1* mutations^11^ as well as DS newborns without *GATA1* mutations^12^, and the risk of acute lymphoblastic leukaemia (ALL) is >30-fold increased in children with DS. Nevertheless, what happens to fetal BM haematopoiesis in DS remains uncertain.

Longer term haematopoiesis depends on a finite pool of haematopoietic stem cells (HSCs), supported by their niche. In mice, BM development begins once vascularization allows HSCs to enter collagenous bone^13^. Stromal cells that support HSC development in humans have been identified but no systematic examination of BM development has been achieved to date. The niche is accepted to be critical to HSC function and niche abnormalities are implicated in both malignant and non-malignant disorders^14^.

In this paper, we use single cell transcriptomics to dissect the composition of human fetal BM in the health and in DS, as haematopoiesis emerges and develops during the early second trimester. We validate: i) newly emerging cell states in fetal BM by FACS-based prospective isolation and single cell RNA sequencing (scRNA-seq) and morphology; and ii) HSC differentiation potential using a single cell clonogenic differentiation assay. We reconstruct haematopoietic differentiation trajectories and predict receptor-ligand interactions to infer the stromal support required for differentiation. Drawing upon existing scRNA-seq data from FL and adult BM, we show for the first time in humans how life-long definitive haematopoiesis is established in the BM early in fetal life and the unique characteristics that contribute towards the assembly of a complex multilineage blood and immune system within a matter of weeks.

## Results

### A single cell transcriptome map of human fetal BM

To characterize both fetal BM haematopoiesis and the environment that supports it, we obtained single cell suspensions from mechanical disruption of fetal femur. Suspensions were FACS sorted into CD45^+^ and CD45^-^ gates, allowing representation of haematopoietic and non-hematopoietic populations for scRNA-seq profiling (10x Genomics) (**Fig. 1A**). Isolates were obtained from 9 karyotypically normal fetuses from the time of macroscopically detectable haematopoiesis (12 PCW) until established 2nd trimester haematopoiesis (18-19 PCW).

**Figure 1:**
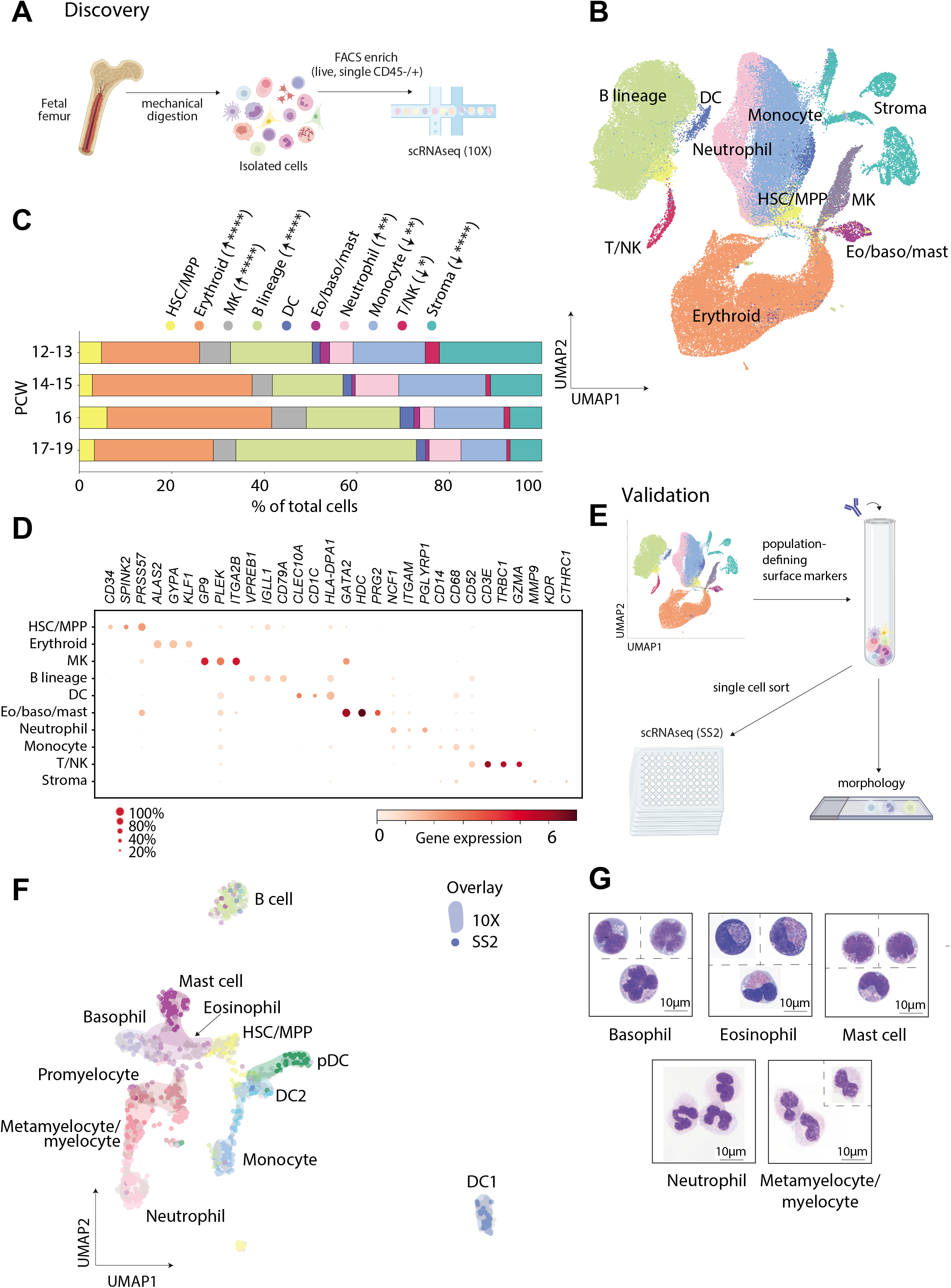
A single cell transcriptome map of human fetal BM. (**A**) Overview of the experimental protocol used to generate scRNA-seq data from fetal BM samples. (**B**) UMAP visualisation of the fetal BM dataset with broad annotation of cell states (= 104,562). (**C**) Bar plot showing broad cell states in fetal BM across gestational age. Statistical significance of cell frequency change by stage shown in parentheses (negative binomial regression with bootstrap correction for sort gates; *\*p* < 0.05, \*\**p* < 0.01, \*\*\**p* < 0.001, \*\*\*\**p* < 0.0001. (**D**) Dot plot showing gene expression of cell-state defining genes. Log-transformed, normalised and scaled gene expression value is represented by the colour of the dot. Percentage of cells in each cell type expressing the marker is shown by the size of the dot. (**E**) Overview of the experimental protocol used to validate cell states by FACS sorting and plate-based scRNA-seq (SS2). 12 cell states were gated using antibodies to markers identified in 10x expression data (**Supplementary Fig. 1**). (**F**) UMAP visualisation of validation SS2 data (= 486) with a 50-cell subset of predicted celltype 10x counterparts (n=600). Cell states in 10x data are represented by coloured areas and cell states in SS2 data are represented by dots of the equivalent colour. (**G**) Cytospin images of selected populations sorted according to gating strategy shown in **Supplementary Fig. 1** and stained with Giemsa. 100x images were concatenated as shown by dotted lines.

Fetal BM 10x data comprised 122,584 cells, of which 104,562 passed quality control (QC) and doublet exclusion (see Methods). Harmony was used to integrate data across batches (**Supplementary Fig. 1**). Data were partitioned by graph-based Leiden clustering, and differentially expressed genes (DEGs) were used to annotate cell states by reference to published literature. Over 60 transcriptionally distinct cell states were identified and manually grouped into 10 broad cellular compartments (**Fig. 1B-D**).

From initiation of haematopoiesis at 12 PCW to establishment at 18-19 PCW, the ratio of haematopoietic: stromal cells expanded rapidly from 3:1 to 13:1 (**Fig. 1C**). B-lymphopoiesis progressively expanded over gestation while the total myeloid proportion remained the same (**Fig. 1C**). A panel of population-defining DEGs expressed as surface antigens was selected to prospectively isolate cell states for validation (**Fig. 1E, Supplementary Fig 1**). Cell states not previously validated in fetal tissues^3,4,15,16^ were prioritized, with emphasis placed on myeloid innate immune cells that were not seen in FL haematopoiesis^4^, particularly granulocytes and their precursors. Cells were FACS-isolated into plates for Smart-seq2 (SS2) analysis and cytospins were prepared for morphological assessment. There was good correlation between 10x and SS2 transcriptional states (**Fig. 1F**) and morphology-based identity assignment (**Fig. 1G**).

Data sets representing haematopoiesis across the human lifespan were assembled for comparison (**Supplementary Fig. 1**). Published, annotated data on fetal YS (4-7 PCW, n=3, k=10,071), FL (7-17 PCW, n=14, k=113,063) and thymus (7 PCW-35 years, n=24,k=259,265) were used^3,4^. Publicly available data on cord blood (CB) (n=4, k=148,442) and adult BM (26-52 years, n=4, k=142,026) (https://data.humancellatlas.org/) were clustered and annotated as above with refined and broad cell type groupings created in line with the fetal BM (**Supplementary Fig. 1**).

### Diversification of innate myeloid and lymphoid cells

Clustering and DEG analysis identified 15 monocyte, DC and neutrophil lineage cell states spanning from committed precursors to terminally-differentiated cells (**Fig. 2A, Supplementary Fig. 2**). These lineages were significantly expanded in fetal BM compared with FL. Fetal BM uniquely contained neutrophilic granulocytes (promyelocytes, myelocytes and neutrophils) (**Fig. 2B**). Additional DC cell states (pDC, tDC and DC3) were observed in fetal BM compared with FL. Fetal BM monocyte, DC and neutrophil diversity was comparable to adult BM with the exception of CD16^+^ monocytes (characterized by *FCGR3A, HES4, CSF1R* co-expression) and monocyte-DC populations.

**Figure 2:**
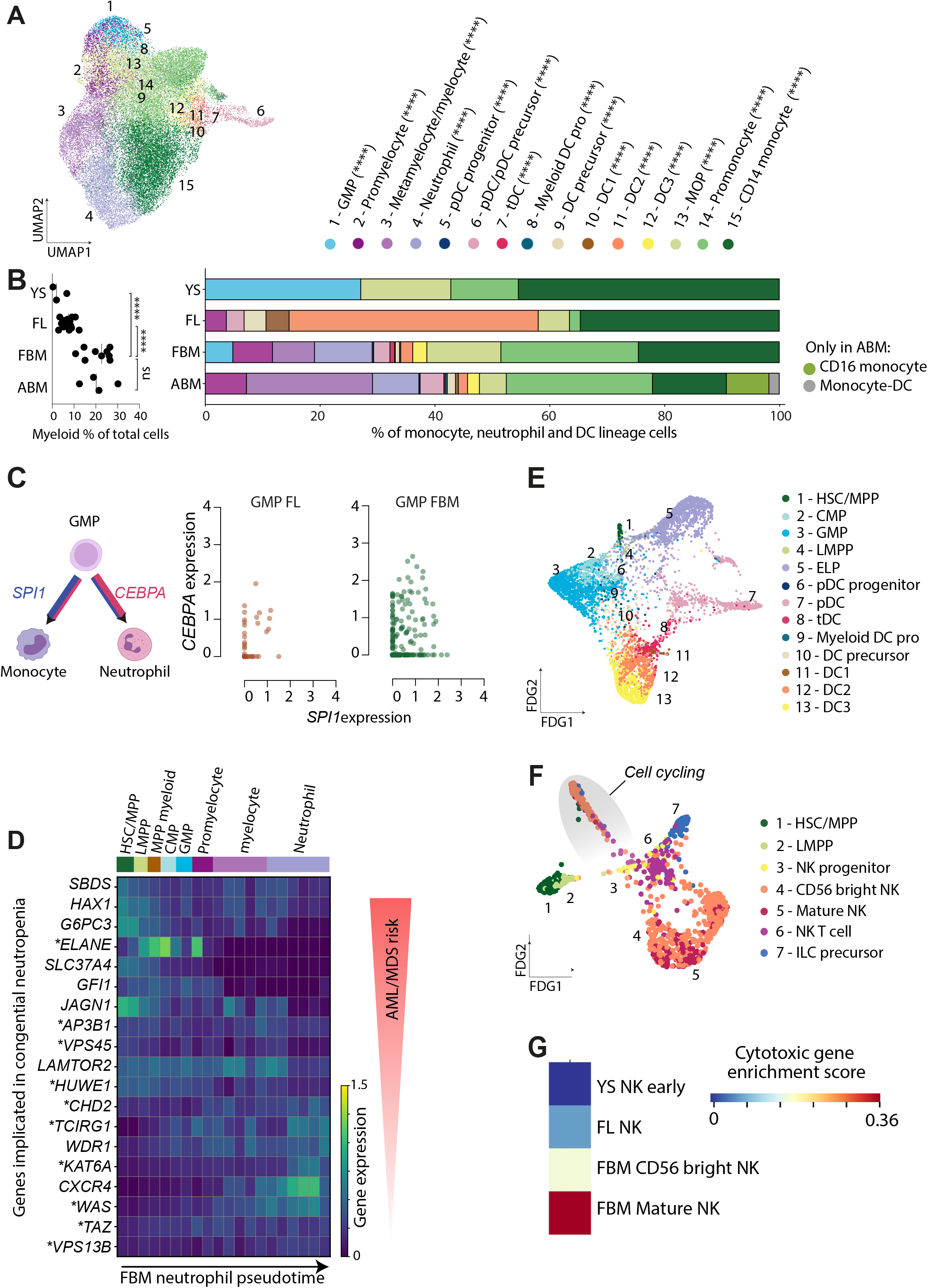
Diversification of innate myeloid and lymphoid cells. (**A**) UMAP visualisation of fetal BM myeloid cells (= 35,008). Cell type is represented by colour, as shown in legend shared with Fig. 2B. (**B**) Left: scatterplot detailing the percentage of myeloid lineage cells present in each sample within YS, FL, fetal BM (FBM) and adult BM (ABM) scRNA-seq datasets (statistical significance tested by one-way ANOVA with Tukey’s multiple comparison tests. Right: bar plot showing the cell states constituting the myeloid lineage within YS, FL, fetal BM and adult BM scRNA-seq datasets. Statistical significance of myeloid cell frequency change by tissue is shown in parentheses (negative binomial regression with bootstrap correction for sort gates; *\*p* < 0.05, \*\**p* < 0.01, \*\*\**p* < 0.001, \*\*\*\**p* < 0.0001). (**C**) Left: Illustration displaying interaction of *SPI1* and *CEBPA* differentiation of monocytes and neutrophils from GMPs. Right: log, normalised and scaled expression of *CEBPA* and *SPI1* in GMP cells from FL (orange) and fetal BM (green), with each dot representing a single cell. (**D**) Heat maps showing expression of genes implicated in severe congenital neutropenia across a Monocle 3-inferred differentiation trajectory of fetal BM neutrophils. Expression values are log-transformed, normalized and scaled. Genes differentially expressed across pseudotime are marked with an asterisk. (**E**) FDG visualisation of fetal BM DC cell states (k = 5,708). (**F**) FDG visualisation of fetal BM NK and ILC cell states (k = 916). Grey ellipse highlights proliferating cells. (**G**) Heat map showing cytotoxicity of NK cell states in YS, FL and fetal BM, as defined by enrichment of genes in the KEGG NK cytotoxicity pathway (See Methods). Relative enrichment is indicated by colour scale.

We sought to explore the origins of neutrophil commitment in fetal BM and identify where FL and fetal BM myeloid differentiation programs diverge. By force-directed graph (FDG), fetal BM progenitor states-HSC, myeloid multipotent progenitor (MPP myeloid) and common myeloid progenitor (CMP)- and the earliest neutrophilic cells (promyelocytes) were bridged by a granulocyte monocyte progenitor (GMP)-like population that was much less prominent in FL^4^ (**Supplementary Fig. 2**). The GMP was upstream of both neutrophils and monocytes by FDG, but multiple lines of evidence indicate that neutrophil versus monocyte fate decisions are made early in myelopoiesis, with the majority of GMPs already committed to either fate^17,18^. Consistent with this, we discovered a cell state that we termed a committed monocyte progenitor (MOP), as it closely resembled the recently described committed monocyte progenitor in CB and adult BM^19^, lying between GMP and promonocytes on the monocyte differentiation trajectory. Prompted by this, we further analysed our previously published FL transcriptomic data and identified the presence of MOP and a small number of promyelocytes within the “neutrophil-myeloid progenitor” population. We generated fetal BM monocyte and neutrophil gene signatures and used these to profile single cells along the continuum of myeloid progenitors, noting that GMPs expressed signatures of one or the other fate (**Supplementary Fig. 2**). Furthermore, as the balance of transcription factors CEPBα (*CEBPA*) and PU.1 (*SPI1*) is critical for neutrophil versus monocyte specification^20^ we examined the relative expression of *CEBPA* and *SPI1* in FL versus fetal BM GMPs (**Fig. 2C**). FBM GMPs expressed higher *CEBPA* relative to *SPI1* than FL GMP.

The trajectory of neutrophil lineage cell states was inferred using Monocle 3 and genes dynamically regulated over pseudotime were identified. We used this to explore the temporal expression of genes proven critical to neutrophil development by their implication in congenital neutropenias. Genes associated with MDS/AML-risk neutropenias (*SBDS, HAX1, G6PC3* tended to be expressed in HSC/MPPs, whereas genes causing congenital neutropenias without recognized MDS/AML-risk (*AP3B1, CXCR4*) were expressed in the terminal stages of differentiation (**Fig. 2D**).

Given the diversity of DCs in adult humans, fetal BM cell states were aligned with those already well-characterized by scRNA-seq in adult blood^23^ (**Supplementary Fig. 2**). DC and monocyte cell states in YS, FL and adult BM were included for comparison. The transcriptional signatures of DC1 and pDCs (DC6) were highly conserved across haematopoietic time and tissue, whereas DC2 and DC3 signatures were more variable. To explore the origins of fetal BM DC diversity, we visualized terminal DC states and their putative precursors by FDG (**Fig. 2E**). This revealed tDC (DC5) as a transcriptional state intermediate between DC2 and pDC. FDG confirmed a dual origin of pDCs, from myeloid and lymphoid precursors, as previously reported in FL^4^. Using iRegulon we found that many of the TFs predicted to drive terminal differentiation of pDC and tDC were shared (**Supplementary Fig. 2**).

We also identified innate lymphocytes, NK cells, NKT-like cells and ILC precursors in fetal BM (**Fig. 2F**). Sub-clustering CD56 bright NK cells, we identified an NK progenitor state with expression of lymphoid lineage progenitor genes (*SOX4, TOX*) (**Supplementary Fig. 2**). FDG was consistent with NK progenitors emerging from HSC via the lymphoid primed multipotent progenitor (LMPP). NK cells and ILCs have been identified by scRNA-seq in FL and fetal skin prior to 12 PCW, suggesting that peripheral tissues are seeded prior to BM NK and ILC precursor production^4^. Fetal BM NK cells expressed genes related to NK cytotoxicity, in contrast to FL and YS NK cells (**Fig. 2G**).

### Establishment of the adaptive immune repertoire

The post-natal and adult mammalian BM is recognized as a critical site for B cell differentiation, positive and negative selection^24^. However, the extent to which fetal BM contributes to the prenatal B cell repertoire is not fully understood^25^. We delineated 5 distinct cell states in the B cell lineage and observed two bursts of proliferative activity, at the Pre pro-B progenitor and the Pre-B precursor stages (**Fig. 3A-B, Supplementary Fig. 3**). Leveraging corresponding 10x BCR-enriched VDJ data from 2 fetal BM samples at 15 PCW, we detected productive heavy chain rearrangement from the Pre-B precursor stage and productive heavy+light chain from the Immature B cell stage (**Fig. 3B**). The emerging B cell repertoire was diverse, with only a small number of shared clonotypes detected in the Pro-B progenitor and proliferating fractions (Pre-pro B progenitor and the cycling branch of Pre-B precursor) (**Supplementary Fig. 3**). The proportion of B progenitors per mononuclear cells in fetal BM was 10-fold higher than in adult BM, indicating that fetal BM B lineage composition was markedly skewed towards the earlier cell states (**Fig. 3C**).

**Figure 3:**
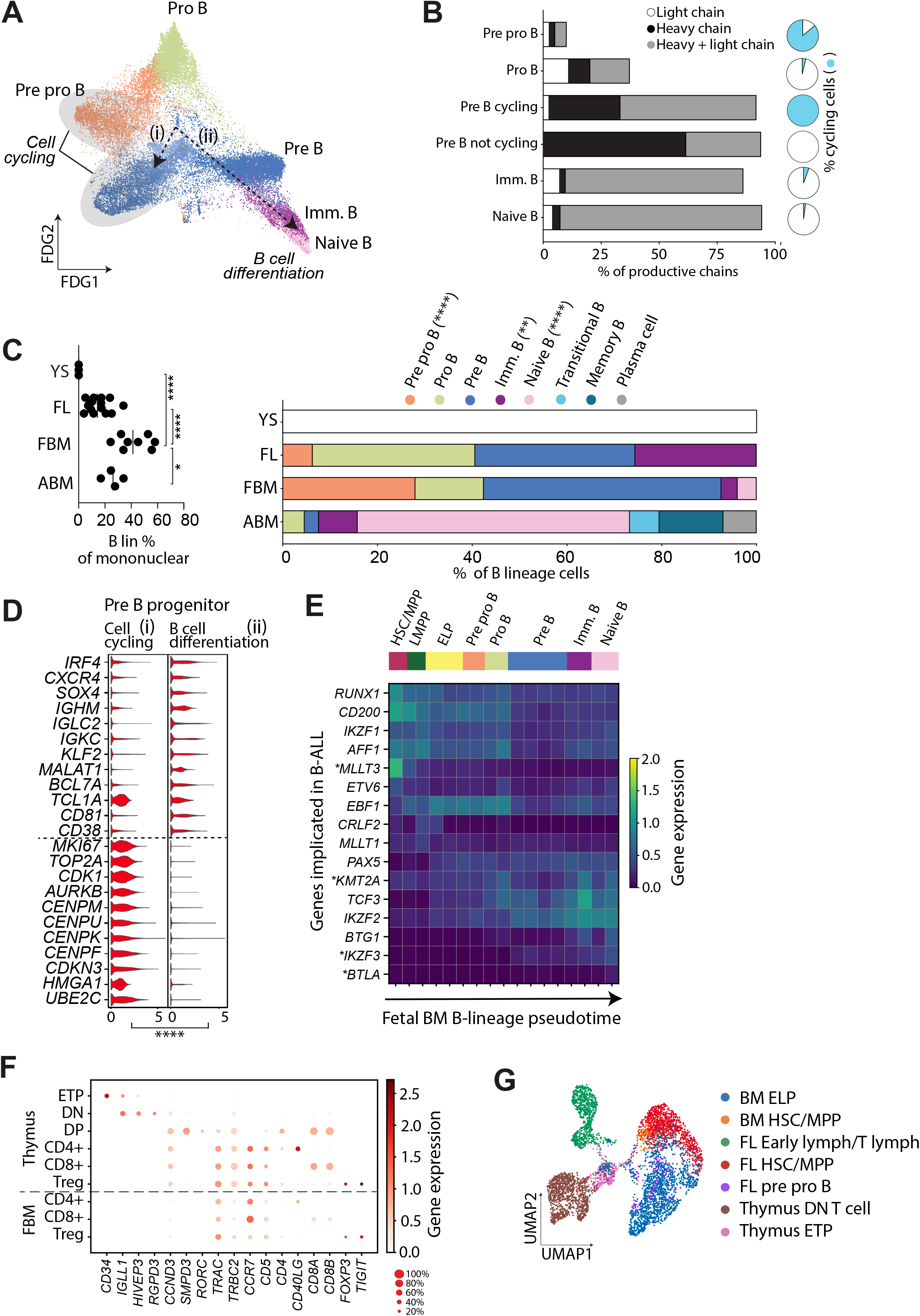
Establishment of the adaptive immune repertoire. (**A**) FDG visualisation of fetal BM B lineage cell states (k = 28,613). Grey ellipses highlight proliferating cells. Pre pro B= Pre Pro B progenitor, Pro B= Pro B progenitor, Pre B= Pre B precursor, Imm. B= Immature B cell. (**B**) Left: bar plot displaying productive chains as a percentage of total chains, and proportions of heavy and light chains in fetal BM B lineage cells (n = 8,298), as defined by BCR-enriched VDJ data. Right: pie charts illustrating the proportion of cycling cells per B-lineage cell-type (defined as > mean cell cycle score) (**C**) Left: scatter plot detailing the percentage of B lineage cells present in each sample in YS, FL, FBM and ABM scRNA-seq datasets (statistical significance tested by oneway ANOVA with Tukey’s multiple comparison tests). Right: bar plot showing cell states constituting the B lineage in YS, FL, FBM and ABM scRNA-seq datasets. Statistical significance of lymphoid cell frequency change by tissue is shown in parentheses (negative binomial regression with bootstrap correction for sort gates; \**p* < 0.05, \*\**p* < 0.01, \*\*\**p* < 0.001, \*\*\*\**p* < 0.0001). (**D**) Violin-plot of DEGs between Pre-B progenitors in paths (i) and (ii) as per **Fig 3A**. Pre-B cells were assigned to a path based on cell cycle enrichment score, as described in methods. Expression values shown on y axis are log-transformed, normalised and scaled. All genes are differentially expressed between both conditions, with *p*<0.0001 (****). (**E**) Heat map showing expression of genes implicated in B-ALL across a Monocle-inferred differentiation trajectory of fetal BM B cells (details in **Supplementary Fig. 3C**). Expression values are log-transformed, normalized and scaled. Genes differentially expressed across pseudotime are marked with an asterisk. (**F**) Dot plot comparing expression of cell state-defining markers genes in fetal thymus and fetal BM (FBM). Dotplot constructed as per Fig. 1D legend and methods. (**G**) UMAP visualisation of fetal BM, FL and thymus progenitors and T precursors (= 4,767). ELP=early lymphocyte precursor, DN T cell= double negative T cell, ETP= early thymocyte precursor.

Differentiation trajectories reconstructed using Monocle revealed linear differentiation from HSC/MPP to ProB progenitors with a subsequent branch-point at the Pre-B cell stage (**Supplementary Fig. 3C)**. One path of Pre-B precursors (“B cell differentiation”) differentiated into immature and naive B cells, significantly expressing TFs and proteins characteristic of B cell maturation e.g. *TCL1A, CXCR4, SOX4* **(Fig. 3D)**. The other Pre-B path (“Cell cycling”) significantly upregulated proliferative genes including *MKI67, TOP2A* and *CDK1*. Apoptosis gene expression was relatively minimal in the “Cell cycling” path, suggesting that cells in this state were capable of returning to the path of progressive differentiation (**Supplementary Fig. 3D**). Apoptosis genes were most enriched in the Pro-B and Pre-B stages, in keeping with programmed death of cells failing successful heavy chain recombination and heavy chain integration into the Pre-B receptor respectively.

Genes differentially regulated during B lineage pseudotemporal development were calculated with Monocle and used as a foundation for exploring the normal expression patterns of genes implicated in childhood B-ALL (**Fig. 3E, Supplementary Fig. 3**). BALL most commonly presents in the first five years of life and leukaemia-initiating events are known to occur *in utero*^8^. Small deletions and translocations in a limited set of genes are well characterized in B-ALL^26^. These genes were highly expressed in fetal BM B cell states, especially in the Pre-B precursor and earlier stages (**Fig. 3E**), while more limited expression was seen in adult BM B cell states (**Supplementary Fig. 3**). As proliferation status and chromatin accessibility contribute to mutagenesis risk^27^, it is possible that the dependence of proliferating B progenitors on these genes exposes a substrate for mutagenesis that is unique to early life haematopoiesis.

While the adult BM is a considerable reservoir of both naive and memory T lymphocytes (**Supplementary Fig. 1**), the fetal BM contains few T lymphocytes (**Fig. 1B**). Thymic lymphopoiesis is established several weeks before fetal BM is colonized^3^, and in keeping with this, we found CD4, CD8 and T regulatory cells expressing genes characteristic of the post thymic state (**Fig. 3F**). TCR-enriched VDJ data on 2 fetal BM samples at 15 PCW confirmed that these single positive T cell subsets contained productive TRA and TRB transcripts (**Supplementary Fig. 3**). In FL, early lymphocytes were transcriptionally similar to the early thymocyte precursor (ETP) in thymus, supporting the hypothesis that FL lymphoid precursors seed the thymus^4^. We combined early lymphoid precursor states from the FL, thymus and fetal BM and identified a proportion of fetal BM early lymphocyte precursors (ELP) with a transcriptional similarity (**Fig. 3G**) suggesting that, in keeping with previous reports, fetal BM contributes to thymopoiesis^28^.

### Intrinsic features of haematopoietic progenitors

Having characterized the committed immune populations of fetal BM, we sought to identify the progenitors responsible for other haematopoietic lineages. We identified 3,746 progenitors, including 92 HSCs expressing *MLLT3, AVP, CLEC9A, HLF* and *HOPX* (**Fig. 4A, Supplementary Fig. 4**). Progenitor populations adjacent to the HSC on the FDG embedding expressed these primitive markers in addition to lineage markers of myeloid (*MPO, CSF1R, CEBPA*) or lymphoid (*BCL11A, IL7R, IL2RG*) fate (**Fig. 4A-B**). Sub-clustering the HSC population did not reveal any further heterogeneity in the HSC compartment or any MPP states with alternative lineage bias. The LMPP was contiguous with the ELP on FDG visualisation, while the myeloid MPP was contiguous with CMP and GMP cell states (**Fig. 4A, Supplementary Fig. 4**).

**Figure 4:**
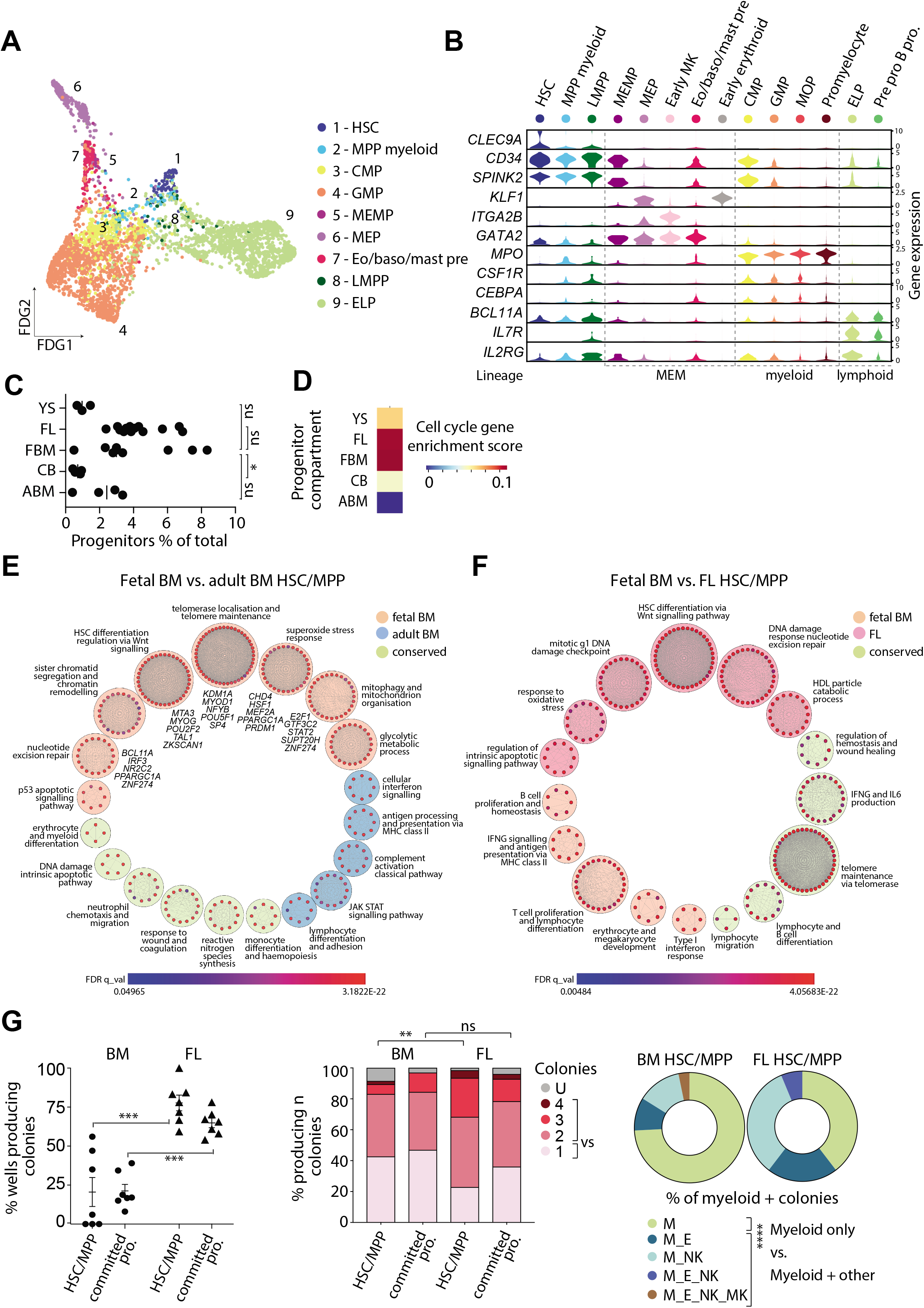
Intrinsic features of haematopoietic progenitors. (**A**) FDG visualisation of fetal BM progenitor cells (= 3,746). (**B**) Violin plots showing expression of progenitor-, MEM-, myeloid- and lymphoid-lineage genes in fetal BM progenitor cells. Log-transformed, normalised and scaled gene expression values are displayed on the y-axis. (**C**) Scatterplot detailing the percentage of progenitor cells present in each sample in YS, FL, fetal BM (FBM) and adult BM (ABM) scRNA-seq datasets (significance tested using a one-way ANOVA with Tukey’s multiple comparison tests). (**D**) Cell cycle gene enrichment score of progenitor compartment cells in YS, FL, fetal BM, cord blood and adult BM. Relative enrichment is indicated by colour scale. Definition of ‘progenitor compartment’ detailed in *Statistics and reproducibility* beneath relevant tissue. (**E**) Chord-plot of conserved and differential pathways in fetal and adult BM HSC/MPPs, derived from Markov Clustering (MCL) of over-represented genesets between analogous cellstates. Large circles represent clusters of functional modules-common to both HSC/MPPs (green), overexpressed in fetal BM HSC/MPPs (orange) and overexpressed in adult BM HSCs (blue). Nodes within each cluster represent significantly enriched gene sets (p <0.05) and edges are weighted by shared genes between gene sets. TFs significantly inferred (p<0.05) to regulate functional modules are listed in italic text. (**F**) Chord-plot of conserved and differential pathways in fetal BM and FL HSCs (method and visualisation as in **Fig 4E**). (**G**) Culture outputs from paired FL and fetal BM HSC/MPP (conducted as per **Supplementary Fig. 4**). Left panel: proportion of sorted wells producing colonies, separated by progenitor type and tissue of origin (n=3 paired fetal BM and FL, n=4 fetal BM only). \*\*\**p*=0.006 by Mann Whitney test of HSC/MPP vs HSC/MPP and \*\*\**p*=0.006 committed progenitor vs committed progenitor. Middle panel: Well contents analysed by flow cytometry and number of lineage outputs per well compared between HSC/MPP and committed progenitors from FL vs. fetal BM. U=colony present but lineage undefinable by this assay. Statistical comparison is of unipotential vs. multipotential colonies: HSC/MPP FL vs fetal BM ** *p*=0.0012; committed progenitor FL vs BM “ns” *p*=0.36 by binomial test. Right panel: Proportion of FL vs fetal BM HSC/MPPs producing myeloid-containing colonies. Statistical comparison is of “myeloid-only” vs. ‘myeloid plus other’ *** *p*=0.0002 by binomial test.

In contrast to FL, which lacked basophils and eosinophils, we found a separate eo/baso/mast precursor in fetal BM expressing *CD34, PRSS57, CSFR2B* and eo/baso/mast lineage genes including *CLC*, similar, at the transcriptomic level, to that reported in adult BM^29^ (**Fig. 4A, Supplementary Fig. 4**). The eo/baso/mast precursor shared gene module expression with megakaryocytes (MK) and erythroid cell states and had transcriptional similarity with the MK-erythroid-mast cell precursor (MEMP). However, it was not possible to infer a developmental relationship between these two progenitors due to scarcity of the MEMP. FDG placed both the eo/baso/mast precursor and the MEMP upstream of differentiated eosinophils, basophils and mast cells (**Supplementary Fig. 4**).

Fetal BM and FL progenitors showed similar levels of proliferation and both were more proliferative than CB and adult BM progenitors, as demonstrated by cell cycle gene enrichment scoring (**Fig. 4D**). We derived and annotated modules of conserved and differential pathways enriched between fetal and adult BM HSC/MPPs by Markov Clustering (MCL) of over-represented genesets (**Fig. 4E**). This demonstrated the relative dependence of fetal BM HSC/MPPs on glycolysis (*TPI1, ENO1, LDHA*), suggested more active response to oxidative (*GPX1, HSPB1*) and replicative stress (*MCM7, HMGB1*) and underscored the importance of Wnt signalling in fetal BM HSC/MPPs. TFs underlying ‘fetalness’ of HSC/MPPs were inferred by modelling the hypergeometric distribution of genes in each pathway module to a TF association matrix derived from the Enrichr database of ENCODE and ChEA consensus TF associations (**Fig. 4E**).

Using the same approach, we found that fetal BM and FL HSC/MPPs shared gene modules for both physiological characteristics, such as telomere maintenance, and differentiation pathways, such as B lymphocyte development (**Fig. 4F**). Fetal BM had greater expression of genes involved in myeloid diversification-genes involved in IFNɣ signalling and antigen presentation via MHC class II (**Fig. 4F**). To establish whether transcriptional differences in FL and fetal BM HSCs corresponded to functionally relevant differentiation potential, we sorted single HSC/MPPs from paired FL and BM suspensions and performed single cell clonal cultures over MS5 mouse stroma with cytokines and growth factors supporting erythroid, megakaryocyte, NK lymphoid, monocyte and neutrophil output (**Fig. 4G, Supplementary Fig. 4**). Notably, the ability to produce B cells was not tested in this assay. More committed populations (CD34^+^CD38^-/mid^ and MLP) were sorted for comparison. Compared to FL HSC/MPPs, fewer fetal BM HSC/MPPs produced colonies. Fetal BM colonies showed greater lineage restriction. Myeloid colonies arose frequently from both FL and fetal BM HSC/MPPs but myeloid-restricted colonies (i.e. no other potential) were typical of fetal BM. The intrinsic myeloid-lineage bias in fetal BM HSC/MPPs may, in part, explain the relative concentration of myeloid cells in fetal BM (**Fig. 2B**).

### Perturbed haematopoiesis in Down syndrome

The constitutional chromosomal anomaly trisomy 21 (Down syndrome; DS) is associated with altered haematopoiesis, including myeloid leukaemia of DS arising from *GATA1* mutations *in utero* and elevated risk of acute leukaemia in early childhood^30,31^. We performed scRNA-seq on 4 DS fetal BM samples (12-13 PCW) in which *GATA1* mutations had been excluded. Of 9,239 cells sequenced, 8,662 passed QC and doublet exclusion (see Methods). Cells were clustered and annotated with refined and broad cell type groupings informed by the disomic (non-DS) fetal BM methodology (**Fig. 5A).**

**Figure 5:**
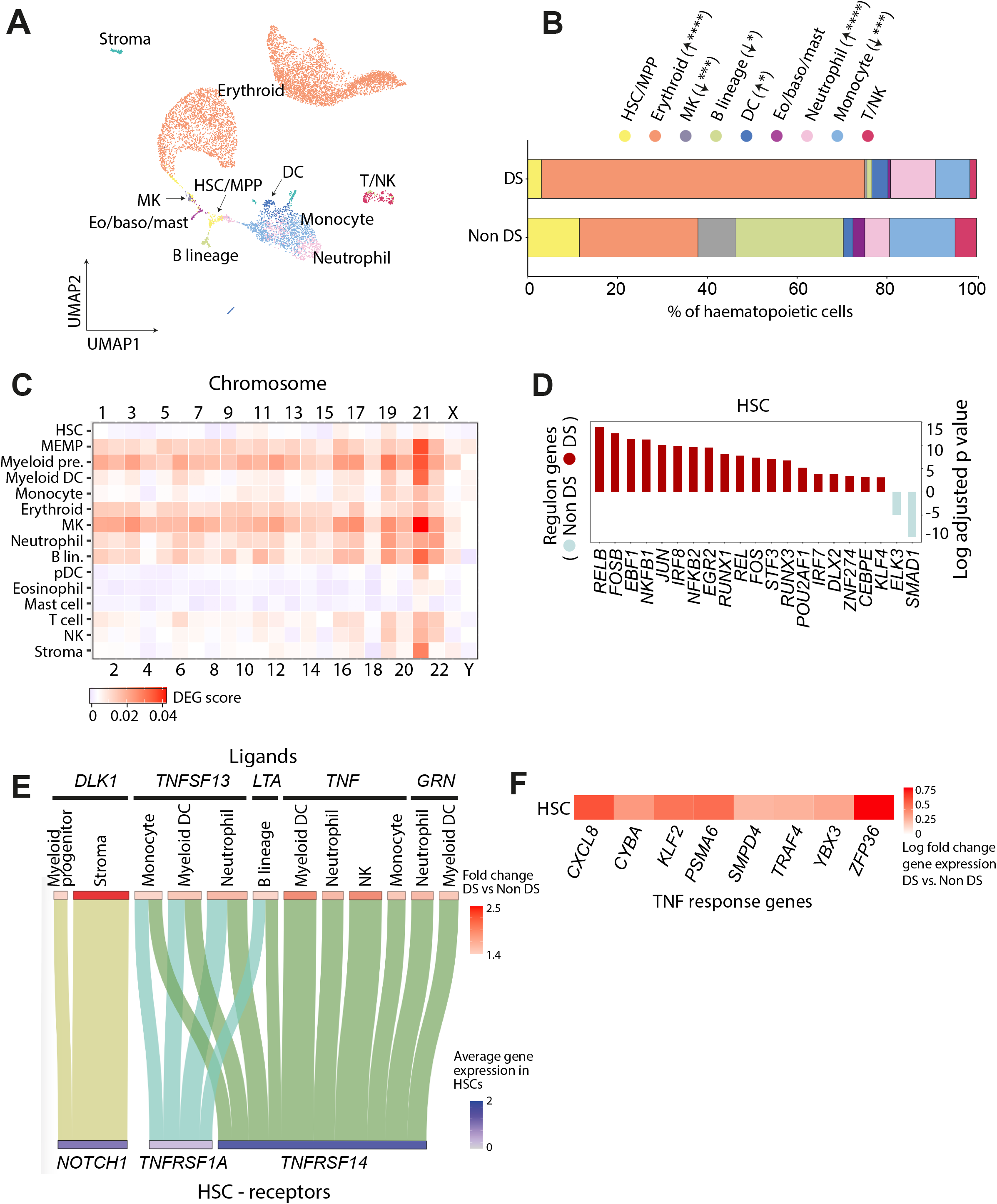
Perturbed haematopoiesis in Down syndrome. (A) UMAP visualisation of fetal BM isolated from 12-13 PCW samples with confirmed trisomy 21 and no *GATA1* mutation (k = 8,662 cells). (B) Stacked barplot showing percentage of cell types present in DS (n=8,662) and non DS fetal BM stage 1 (12 - 13 PCW with n = 10,437). Statistical significance of cell frequency difference is shown in parentheses (negative binomial regression with bootstrap correction for sort gates; \**p* < 0.05, \*\**p* < 0.01, \*\*\**p* < 0.001, \*\*\*\**p* < 0.0001). (C) Heatmap showing number of DEGs between paired cell states in DS vs. non-DS fetal BM for each chromosome, including correction for number of genes per chromosome. (D) Predicted most highly active TFs in DS vs. non-DS HSCs. PySCENIC was used to predict TF activity in each cell. This was used to produce median TF activity in each cluster and DS vs non-DS were compared by t-test for each TF to calculate p-values. Top 20 most significantly differentially active TFs by p-value shown. (E) Sankey plot of putative enriched interactions in DS fetal BM. For each cell category, DEGs between DS and non-DS were filtered for ligands and their receptors were identified using CellPhoneDB. Fold change in expression in DS over non-DS is shown both in the boxes above (red scale) and represented in the thickness of the line between clusters. Below is the predicted receptor coloured by its gene expression level in HSCs (blue scale). (F) Heatmap of differentially expressed TNF response genes by log fold change expression in DS vs. non-DS HSCs. DEGs were filtered for an intersection with TNF response genes from the GO biological process database (GO:0034612).

DS fetal BM had a significantly expanded erythroid compartment and diminished B lymphocyte and MK compartments compared with non-DS fetal BM (**Fig. 5B**). Excluding erythroid cells from analysis, the proportions of B lymphocyte and MK lineages remained significantly lower in DS (**Supplementary Fig. 5**). Relative suppression of the MK lineage in DS fetal BM was not expected as DS FL HSCs show intrinsic erythroid/MK bias^11^. Maturation of erythroid cells was skewed in DS, with an approximately 1:1 ratio of early/mid: late erythroid cells, compared with a 5:1 ratio in non-DS fetal BM (**Supplementary Fig. 5).** Early/mid erythroid cells in DS had higher cell cycle gene enrichment scores. The DC compartment was relatively expanded in DS and probing the composition revealed that the proportion of myeloid DCs (DC1, DC2, DC3) was increased (**Supplementary Fig. 5**).

Erythroid differentiation was reconstructed using Monocle3 in both DS and non-DS fetal BM (**Supplementary Fig. 5**. Genes differentially expressed during erythroid development showed comparable regulation over DS and non-DS erythroid pseudotime (**Supplementary Fig. 5**).

We sought to explore the lineage biases imposed by trisomy 21 on HSCs as the progenitor common to MK, erythroid and B lineage cells. DEGs were calculated between paired cell-states in DS and non-DS fetal BM. While many DEGs were located on chromosome 21, genome-wide transcriptional differences were observed (**Fig. 5C**). In addition, DEGs were noted in B lineage cells, myeloid progenitors, neutrophils and DS stroma (**Fig. 5C**), showing the impact of trisomy 21 extends beyond a single chromosome and beyond haematopoietic progenitors. Myeloid lineages (neutrophils, monocytes, myeloid DCs) had the most DEGs between DS and non-DS fetal BM (**Fig. 5C**).

The observation that DS stromal cells had an altered gene expression profile (**Fig. 5C**) led us to question whether extrinsic dysregulation of haematopoiesis may occur in DS. We filtered DEGs between DS and non-DS for equivalent cell types, selecting for ligands, and used CellPhoneDB to generate predictions of receptor-ligand interactions. We found TNF family ligands (*TNF, LTA, TNFSF13*) and the EGF-like family ligand *DLK1* were overexpressed in DS, particularly in the myeloid lineage (**Fig. 5E, Supplementary Fig. 5**). Ligands enabling physical proximity between HSC and myeloid/ innate lymphoid lineages (SELL, ICAM1) were also expressed more highly in DS (**Supplementary Fig. 5**). DEGs between DS and non-DS HSCs confirmed overexpression of TNF-response genes in DS (**Fig. 5F**).

### Haematopoietic microenvironment in fetal BM

The stromal fraction of fetal BM comprised 6,726 cells. Here we included cells predicted to provide structural or supportive roles, including some cells with haematopoietic origins e.g. macrophages. Fine-resolution clustering revealed 19 cell states (**Fig. 6A, Supplementary Fig. 6**). A number of cell states varied in frequency over gestational age, for example osteoclasts were over-represented in earlier stages and osteoblasts in later stages. Annotation was harmonized with stromal cell states recently described in mouse BM isolated without pre-enrichment^15^ (**Supplementary Fig. 6**). Our most numerous cell states (osteoclasts and macrophages) had no parallel in mouse BM. CXCL12 abundant reticular cells (CAR) in mouse BM had two distinct fractions (osteo-CAR and adipo-CAR) but human fetal BM contained only adipo-CAR. Endothelial cells (EC), osteochondral cells, fibroblasts and Schwann cells correlated well with mouse counterparts (**Supplementary Fig. 6**). We focused our investigation on EC and osteochondral cells which are reported to form the endothelial and endosteal HSC niches^33^.

**Figure 6:**
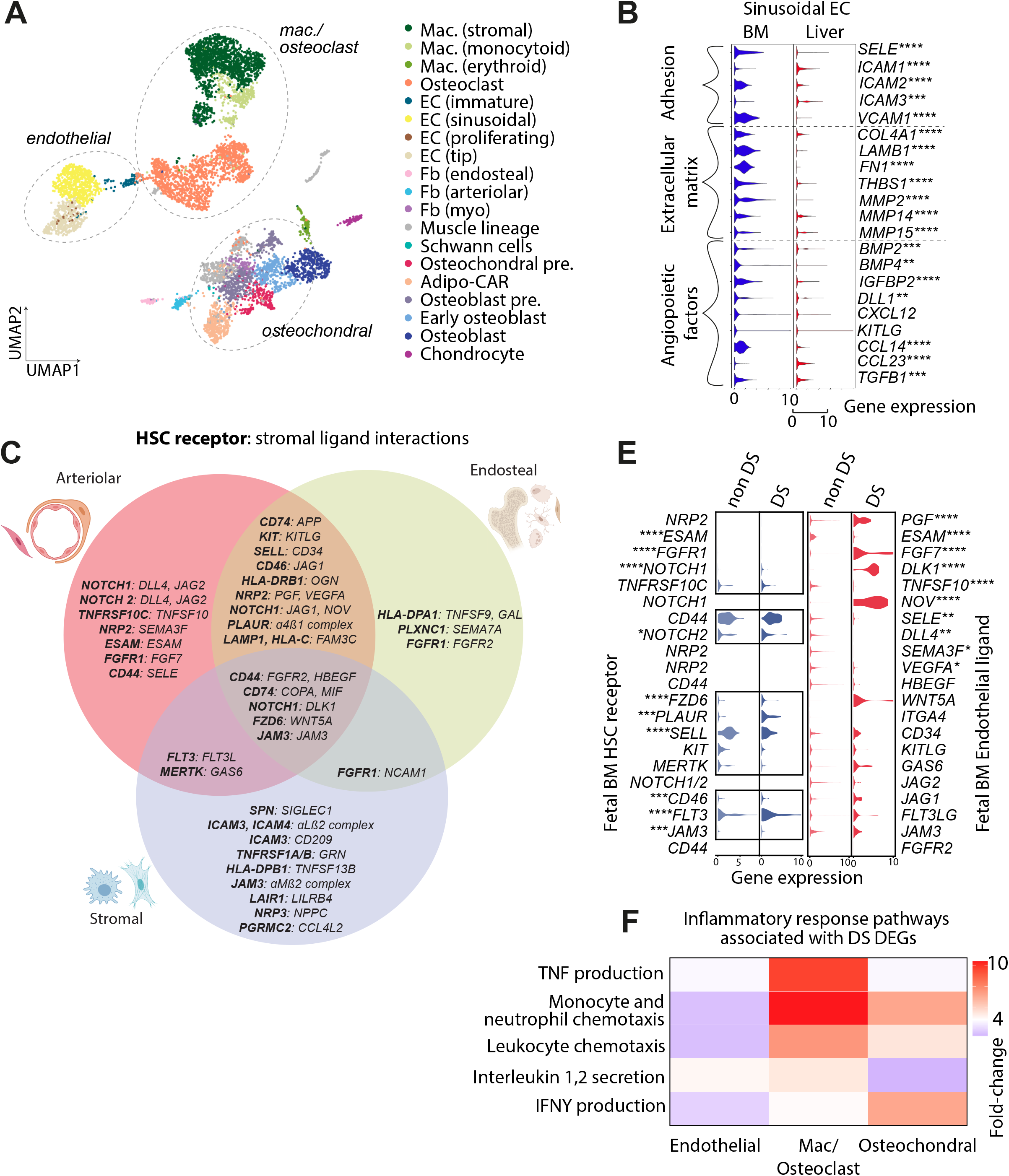
Haematopoietic microenvironment in fetal BM. (**A**) UMAP visualisation of fetal BM stromal cells (k = 6,726). Cell type is represented by colour, as shown in legend. Broad groupings of macrophage/osteoclast, endothelial and osteochondral cell types are shown using dotted lines. (**B**) Violin plot of gene expression in FL and fetal BM sinusoidal EC. Genes with documented role in adhesion, extracellular matrix formation and angiopoiesis are shown. Genes that are differentially expressed between tissues are highlighted with an asterisk (subject to Wilcoxon rank-sum test with Benjamini-hochberg correction for multiple testing; \**p* < 0.05, \*\**p* < 0.01, \*\*\**p* < 0.001, \*\*\*\**p* < 0.0001). (**C**) Summary of receptor-ligand interactions predicted between fetal BM HSC and stromal cells by CellPhoneDB. Cells are grouped into arteriolar (EC-sinusoidal, EC-proliferating, EC-tip, EC-arteriolar, Fb-arteriolar, adipo-CAR), endosteal (Fb-endosteal, Osteochondral precursor, Early osteoblast) and stromal (Mac-stromal, Fb-fibroblast). Significant putative HSC receptor/stromal ligand interactions between are indicated in the Venn diagram. (**D**) Violin plot of gene expression in DS and non-DS fetal BM HSC and ECs. Genes shown have a significant receptor-ligand interaction in non-DS fetal BM, as predicted by CellPhoneDB analysis and detailed in **Fig. 6C**. Genes that are differentially expressed between DS and non-DS fetal BM are highlighted with an asterisk (subject to Wilcoxon rank-sum test with Benjamini-hochberg correction for multiple testing; \**p* < 0.05, \*\**p* < 0.01, \*\*\**p* < 0.001, \*\*\*\**p* < 0.0001). (**E**) Heatmap of differentially enriched inflammatory and cytokine production pathways in DS vs. non-DS stromal cell groups. See methods for further details.

We annotated one cluster with 191 cells as osteochondral precursor as it expressed marker genes for all three osteochondral populations: osteoblasts (*CPE, IBSP*), Adipo-CAR (*VCAN, GGT5*) and chondrocytes (*CSPG4, EDIL3*) in addition to genes with a role in musculoskeletal development (*AEBP1, FOXC1*). Visualized by FDG, the osteochondral precursor was embedded at a central point between the three emerging lineages (**Supplementary Fig. 6**).

Sinusoidal ECs expressing *STAB1, STAB2* and *TSPAN7* were the most abundant EC type in BM (**Fig. 6A, Supplementary Fig. 6**). The sinusoids form a portal of entry and exit for migratory progenitors and differentiated cell types in BM as well as other organs capable of haematopoiesis (liver and spleen). We compared expression of genes between sinusoidal EC in FL and fetal BM to identify factors allowing access to each niche (**Fig. 6B)**. Fetal BM sinusoidal ECs had significantly higher expression of *SELE, VCAM1* and *ICAM2*. Fetal BM sinusoidal ECs expressed more extracellular matrix genes, including *THBS1*, which may contribute to HSC/MPP retention^34^ and matrix metalloproteinases which are reported to facilitate mature cell egress^35^ (**Fig. 6B)**. Both FL and fetal BM sinusoidal ECs expressed genes for secreted factors: fetal BM sinusoidal ECs expressed more *CCL14*, promoting proliferation of myeloid progenitors, while FL expressed more *CCL23* which has the opposite effect^36^, supporting a role for ECs in shaping tissue-specific myelopoiesis.

We used CellphoneDB to predict significant receptor-ligand interactions between fetal BM stromal cells and HSCs (**Fig. 6C, Supplementary Fig. 6**). Interactions with significant association and receptor-ligand expression were grouped by i) directionality of signalling (whether to or from HSC) and ii) niche environment (whether arteriolar, endosteal or stromal; see Methods for niche groupings). EC and adipo-CAR cells were predicted to provide critical signals to support haematopoiesis via CD44, KIT, FLT3, NOTCH1/2, NRP2, MERTK, FZD6 and FGFR1 and to support HSC homing and retention via SELL, JAM3 and ESAM (**Fig. 6C, Supplementary Fig. 6**). Osteochondral cells and stromal cells appeared capable of providing similar support, though NOTCH signalling interactions were fewer (**Fig. 6C, Supplementary Data Fig. 6**).

Stromal cells isolated from DS fetal BM were few in number, precluding the detailed annotation of cell states we performed for non-DS fetal BM. However, EC were readily identifiable and the key interactions predicted between EC and HSC in non-DS fetal BM were examined in DS fetal BM (**Fig. 6E**). NOTCH ligands *NOV, DLL4* and *DLK1* were more abundant in DS endothelium. NOTCH signalling has a critical role in HSC emergence, maintenance and response to proinflammatory signals^37,38^. We probed the relative frequency of proinflammatory transcriptional programmes in DEGs between three broad non-haematopoietic groupings in DS and non-DS (endothelial, macrophage/osteoclast, and osteochondral). TNF production was concentrated in macrophage/ osteoclasts and IFNγ production in osteochondral cells (**Fig. 6F**). Our collective findings reveal an altered stromal environment in DS.

## Discussion

Survival of the fetus depends on successful initiation of haematopoiesis in several organs across gestation. We reveal the complete establishment of haematopoiesis in the FBM within the first few weeks of the second trimester and identify the BM as a key site of neutrophil emergence, myeloid diversification and B lymphoid selection. We identify a unique intrinsic molecular profile of FBM HSC/MPPs, an intrinsic bias of DS BM stem/progenitors underpinned by genome-wide transcriptional changes. A better understanding of human developmental haematopoiesis has the potential to inform regenerative and transplantation therapies, for example, through co-opting developmental programmes to accelerate reconstitution of haematopoietic stem cell transplants, and manipulating the lineage bias of differentiating progenitors to address specific deficiencies or for cellular therapy. For such endeavours to be successful, an initial phase of discovery science is critical. It is in this context that the current study provides the first comprehensive analysis of human FBM haematopoiesis to address a major previous knowledge gap.

## Methods and Supplementary Figures

See Supplementary Information.

## Data and materials availability

The raw 10x fetal BM sequencing data is deposited at ArrayExpress, with accession code E-MTAB-9389.

## Code availability

Single-cell sequencing data were processed and analysed using publicly available software packages. Python/R code and notebooks for reproducing single-cell analyses are available at https://github.com/haniffalab/FCA_bone_marrow

## Acknowledgements

We thank the Newcastle University Flow Cytometry Core Facility, Bioimaging Core Facility, Genomics Facility, NUIT for technical assistance, School of Computing for access to the High-Performance Computing Cluster, CellGenIT, Newcastle Molecular Pathology Node Proximity Lab, and Alison Farnworth for clinical liaison. The human embryonic and fetal material was provided by the Joint MRC / Wellcome (MR/R006237/1) Human Developmental Biology Resource (www.hdbr.org).

## Author Contributions

M.H.; S.A.T; I.R. and B.G. conceived and directed the study. M.H.; S.A.T.; I.R.; B.G.; L.J.; and E.L. designed the experiments and data analysis approach. Samples were isolated by S.L.; R.A.B.; I.G.; J.E.; P.B.; K.A.; S.O.B.; N.E.; libraries prepared by E.P.; and E.S; and sequencing by J.C.; R.Q.; R.H.; and WSI core facility. Flow cytometry and FACS experiments were performed by R.A.B.; L.J. and D.M., supported by D.McD.; and A.F. Cytospins were performed by L.J. and D.D.; and *in vitro* culture differentiation experiments were performed by L.J.; C.M and D.M. Immunofluorescence microscopy was performed by C.J.; T.N.; R.C.; C.C.;. C.S.;. M.A., with analysis performed by M.M., B.O., C.S.; B.P.; and I.G. M.S.K.; B.L.; O.A.; M.T.; D.D.; T.L.T.; M.S.; O.R-R. and A.R. generated adult and cord blood scRNA-seq datasets. CITE-seq datasets were generated by E.S.; N.M.; and N.K.W. Computational analysis was performed by S.W.; I.G.; M.Q.L.; G.R.; E.D.; I.K.; M.M.; J.B.; M.S.J.; M.E.; and web portals were constructed by I.G.; D.H.; and J.McG., with disease information assembled by K.P. and T.C. M.H.; L.J.; S.W.; I.G. G.R.; B.O.; H.K.; K.B.M.; T.C.; N.M.; N.K.W.; K.B.M.; D.H.; D.M.P.; S.B.; A.R.; E.L.; B.G.; I.R.; and I.G.; and S.A.T. interpreted the data. M.H.; L.J.; S.W.; I.G.; G.R.; B.G.; I.R. and S.A.T. wrote the manuscript, with input from M.L.R.H and J.E.L. All authors read and accepted the manuscript.

