## Supplementary Figures 1-6 and Methods for "Intrinsic and extrinsic regulation of human fetal bone marrow haematopoiesis and perturbations in Down syndrome"

### Supplementary Figure 1

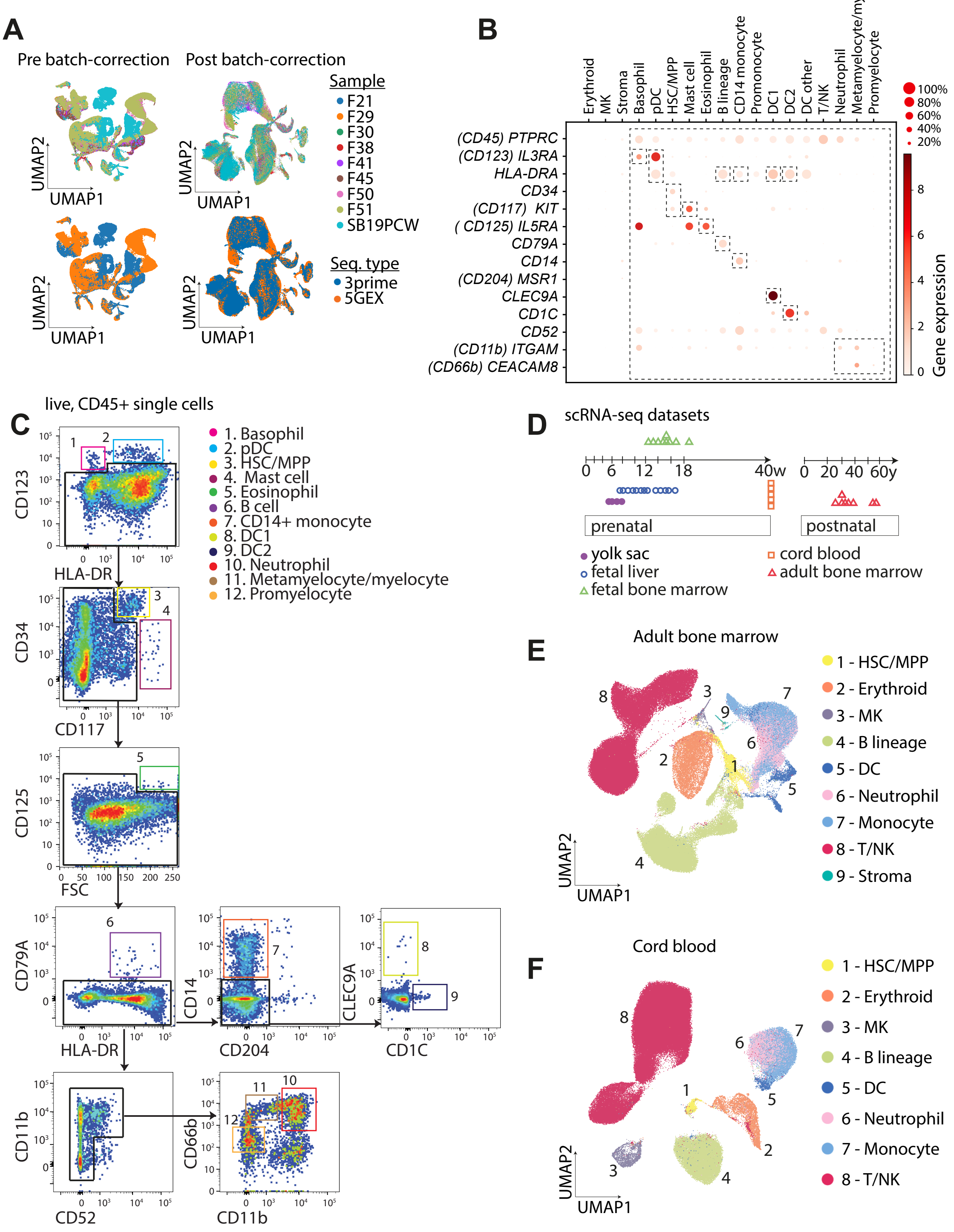

**Supplementary Figure 1: A single cell transcriptome map of human fetal BM**

(**A**) UMAP visualisation of fetal BM scRNA-seq data pre and post batch correction. Sequencing type and sample is represented by colour, as shown in legend.

(**B**) Dot plot showing gene expression of cell-state defining genes in 10x data. Equivalent surface antigen names are shown in parentheses. Dotplot constructed as per Fig. 1D legend and methods. Dashed boxes indicate the gating strategy for cell-sorting, for example pDCs were isolated as CD45$^{+}$HLA-DR$^{+}$CD123$^{+}$ cells.

(**C**) Sorting strategy used to isolate cell types for validation based on subset defining markers from scRNA-seq (10x) data. From live, CD45$^{+}$ single cells, CD123$^{+}$HLA-DR$^{-}$ basophils and CD123$^{+}$HLA-DR$^{+}$ pDCs were gated. From the remaining cells, CD34$^{+}$CD117$^{mid\text{-}hi}$ progenitors and CD117$^{hi}$ mast cells were gated. Next, CD125$^{+}$FSC$^{hi}$ eosinophils were gated. Subsequently, HLA-DR$^{+}$CD79a$^{+}$ B cells were separated. From HLA-DR$^{+}$ cells, CD14$^{+}$CD204$^{-}$ monocytes were gated. Within the CD14$^{-}$CD204$^{-}$ population, CLEC9A$^{+}$ DC1 and CD1c$^{+}$ DC2 were identified. From HLA-DR$^{-}$ cells, CD11b$^{-}$CD52$^{+}$ T and NK cells were excluded. From the remaining cells CD11b$^{-}$CD66b$^{-}$ promyelocytes, CD11b$^{-}$CD66b$^{+}$ metamyelocytes/myelocytes and CD11b$^{+}$CD66b$^{+}$ neutrophils were selected. Note, the CD11b gating was as per published reports mature neutrophils being CD11b$^{+}$ and immature neutrophils being CD11b$^{-}$[42].

(**D**) Summary of scRNA-seq datasets used for comparison: published YS and FL data[4] and publicly available cord blood and adult BM data from the Human Cell Atlas Data Coordination Portal.

(**E**) UMAP visualisation of adult BM scRNA-seq dataset with broad annotation of cell states applied (k = 142,026).

(**F**) UMAP visualisation of cord blood scRNA-seq dataset with broad annotation of cell states applied (k = 148,442).

### Supplementary Figure 2

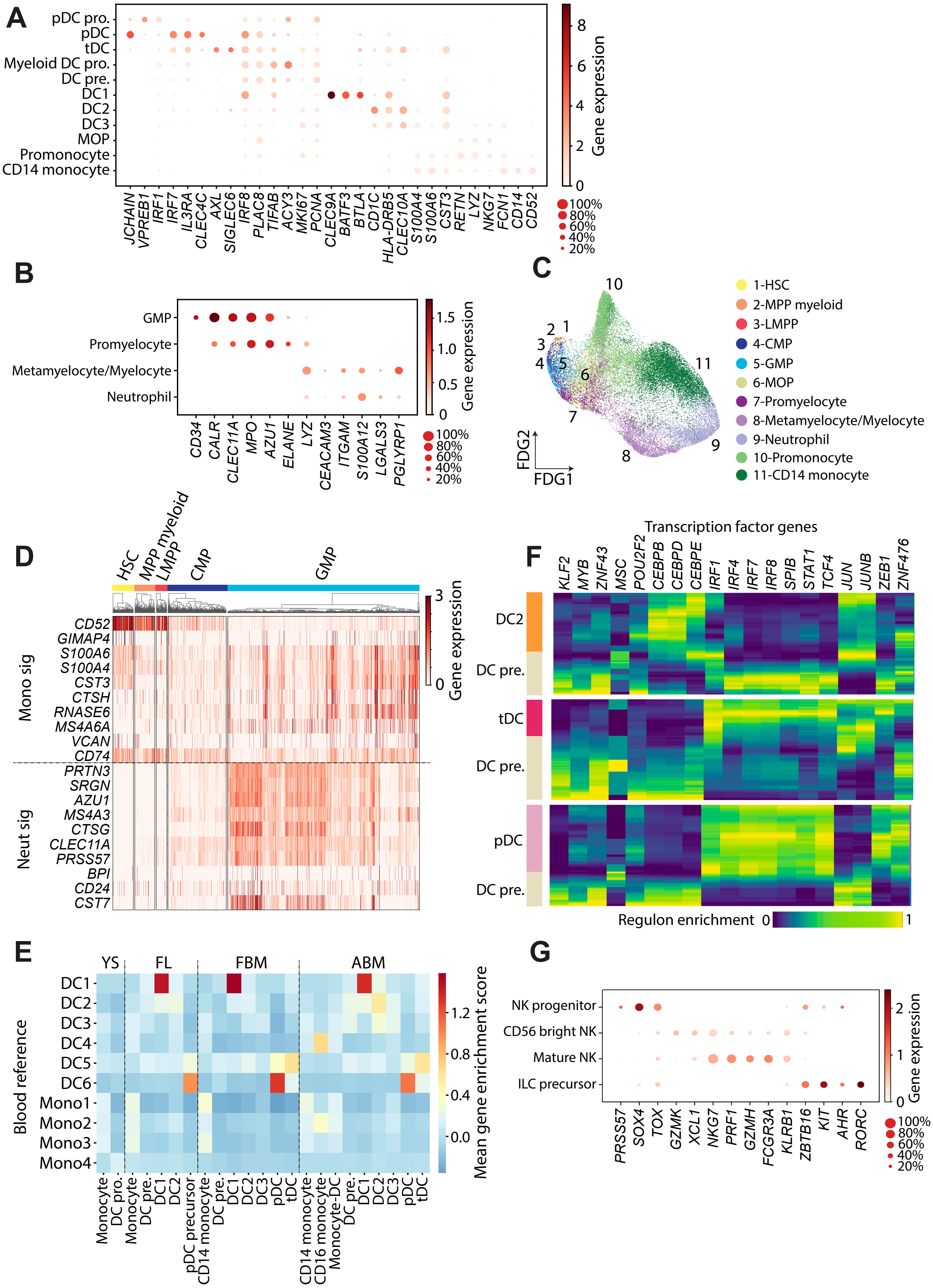

**Supplementary Figure 2: Diversification of innate myeloid and lymphoid cells**

(**A**) Dot plot showing expression of cell state-defining marker genes in fetal BM DC and monocyte lineage cells. Dotplot constructed as per Fig. 1D legend and methods.

(**B**) Dot plot showing expression of cell state-defining marker genes in fetal BM myeloid precursors and neutrophil lineage cells.

(**C**) FDG visualisation of fetal BM progenitor, neutrophil and monocyte cell states (k = 33,137).

(**D**) Heatmap showing gene expression of markers for early monocyte and neutrophil commitment (derived from promonocyte vs. promyelocyte DEGs) in fetal BM progenitor cell states. Gene expression shown is log-transformed, normalised and scaled.

(**E**) Heat map showing transcriptional similarity between DC/monocyte cell states established in blood[23] with those identified in developing and mature haematopoietic tissues (YS, FL [4] fetal BM, adult BM). Gene enrichment scores are based on gene signatures pooled from top 100 statistically significant monocyte/DC DEGs in healthy blood[23].

(**F**) Heatmap of predicted activity of TFs across pseudotime. TF activity was inferred using PySCENIC and pseudotime was calculated in Scanpy (*sc.tl.dpt*). For each TF the expression data was normalised to between 0-1 prior to plotting.

(**G**) Dot plot showing expression of genes in fetal BM NK and ILC cells.

#### Supplementary Figure 3

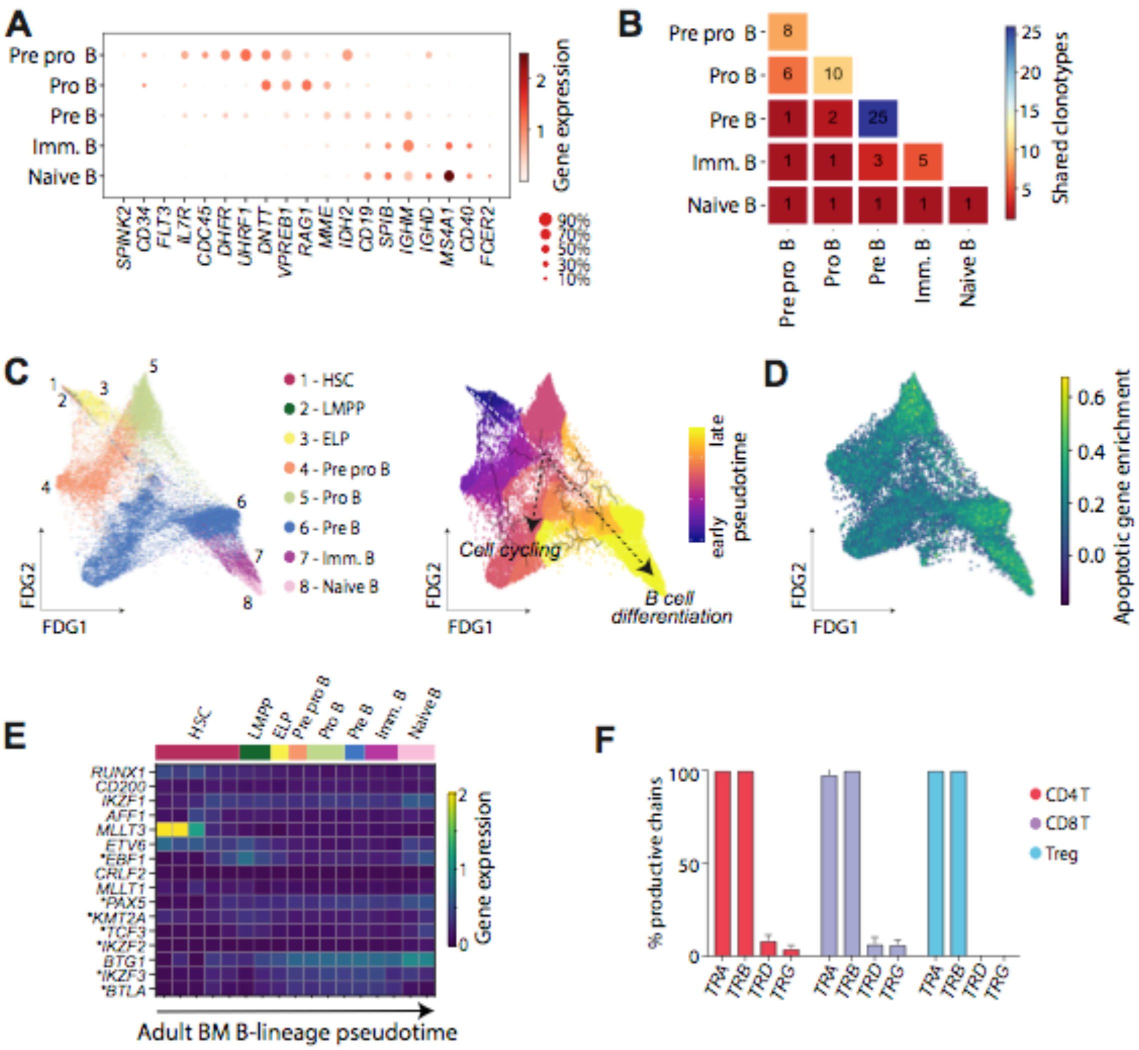

**Supplementary Figure 3: Establishment of the adaptive immune repertoire**

(A) Dot plot showing expression of cell state-defining marker genes in fetal BM B lineage cells. Dotplot constructed as per Fig. 1D legend and methods.

(B) Heatmap showing number of shared clonotypes between B lineage cell types, as defined by CellRanger. Number of shared clonotypes is shown by colour of square, with colour scale in legend.

(C) Left: FDG visualisation of fetal BM B lineage cell states (k = 30,097). Cell state is represented by colour, as shown in legend. Right: FDG visualisation of fetal BM B lineage cell states (n = 30,097). Monocle-inferred pseudotime trajectory is overlaid onto FDG and cells are coloured by pseudotime value, as shown in legend. Two paths inferred from Monocle pseudotime trajectory are depicted by dashed arrows.

(D) FDG visualisation of fetal BM B lineage cell states (k = 28,613). Cells are coloured by apoptotic gene enrichment score, as defined by their expression of genes in the KEGG apoptotic pathway (details in methods).

(E) Heat map showing expression of genes implicated in B-ALL across a Monocle-inferred differentiation trajectory of adult BM B cells. Expression values are log-transformed, normalized and scaled. Genes differentially expressed across pseudotime are marked with an asterisk.

(F) Barplot showing proportion of TRA/B/G/D chains per productively rearranged TCRs by fetal BM T lineage cell state, according to CellRanger VDJ output. Bars (mean) and error bars (SD) of n=2 15 PCW fetal BM samples are shown. Mean±SD percent productivity of TCRs was 93±9%, 81±16% and 92±11% for CD4 T cells, CD8 T cells and Treg.

### Supplementary Figure 4

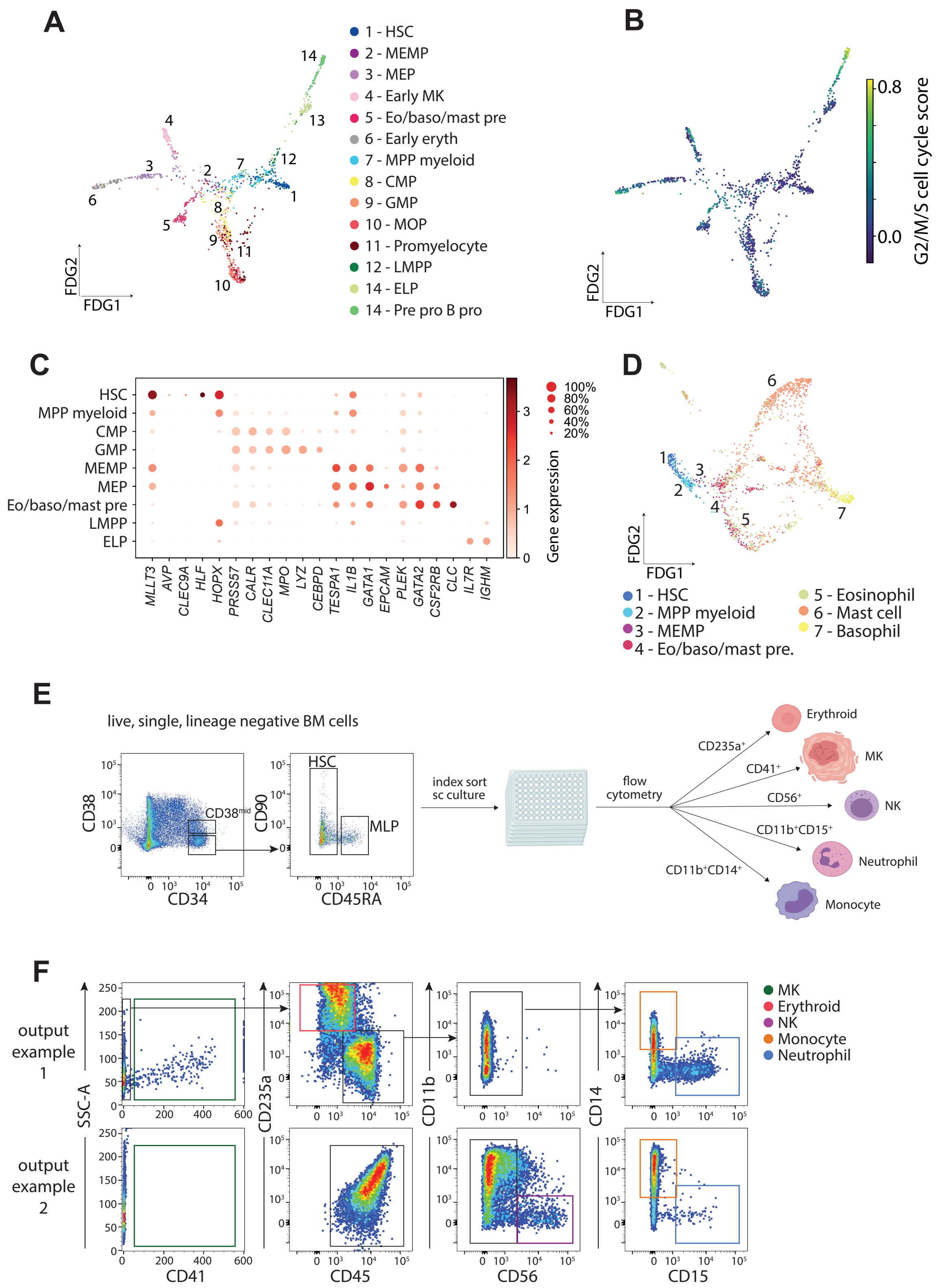

**Supplementary Figure 4: Intrinsic features of haematopoietic progenitors**

(A) FDG visualisation of fetal BM progenitor cells, with maturing cell states down-sampled to allow visual inspection of lineages emerging (k = 934). Observed proportions of cell states are given in **Fig. 4A**.

(B) FDG visualisation of fetal BM progenitor cells, as shown in **Supplementary Fig. 4A,** with colour representing cell cycle gene enrichment score (see scale in legend).

(C) Dot plot showing expression of cell state-defining marker genes in fetal BM progenitor cells. Dotplot constructed as per Fig. 1D legend and methods.

(D) FDG visualisation of fetal BM non-neutrophilic granulocyte cells (k = 1,487).

(E) Sort gates for HSC culture experiments. Single, live, lineage negative $CD34^+$ cells were divided into $CD34^+CD38^{hi}$ (top 20%), $CD34^+CD38^{mid}$ (middle 60%), $CD34^+CD38^-$ (bottom 20%). $CD34^+CD38^-$ cells were gated further into $CD45RA^-$ HSC/MPP and CD45RA LMPP/MLP. HSC, LMPP/MLP and $CD34^+CD38^{mid}$ cells were index sorted for single cell culture on an MS5 stromal layer. LMPP/MLP and $CD34^+CD38^{mid}$ cells were analysed as "committed progenitor".

(F) Examples of HSC culture outputs, each gated on single cells. Example 1 showing $CD41^+$ MK, $CD235a^+$ erythroid, $CD14^+$ monocyte and $CD15^+$ neutrophil outputs. Example 2 showing $CD56^+$ NK, $CD14^+$ monocyte and $CD15^+$ neutrophil outputs (for scoring see methods).

### Supplementary Figure 5

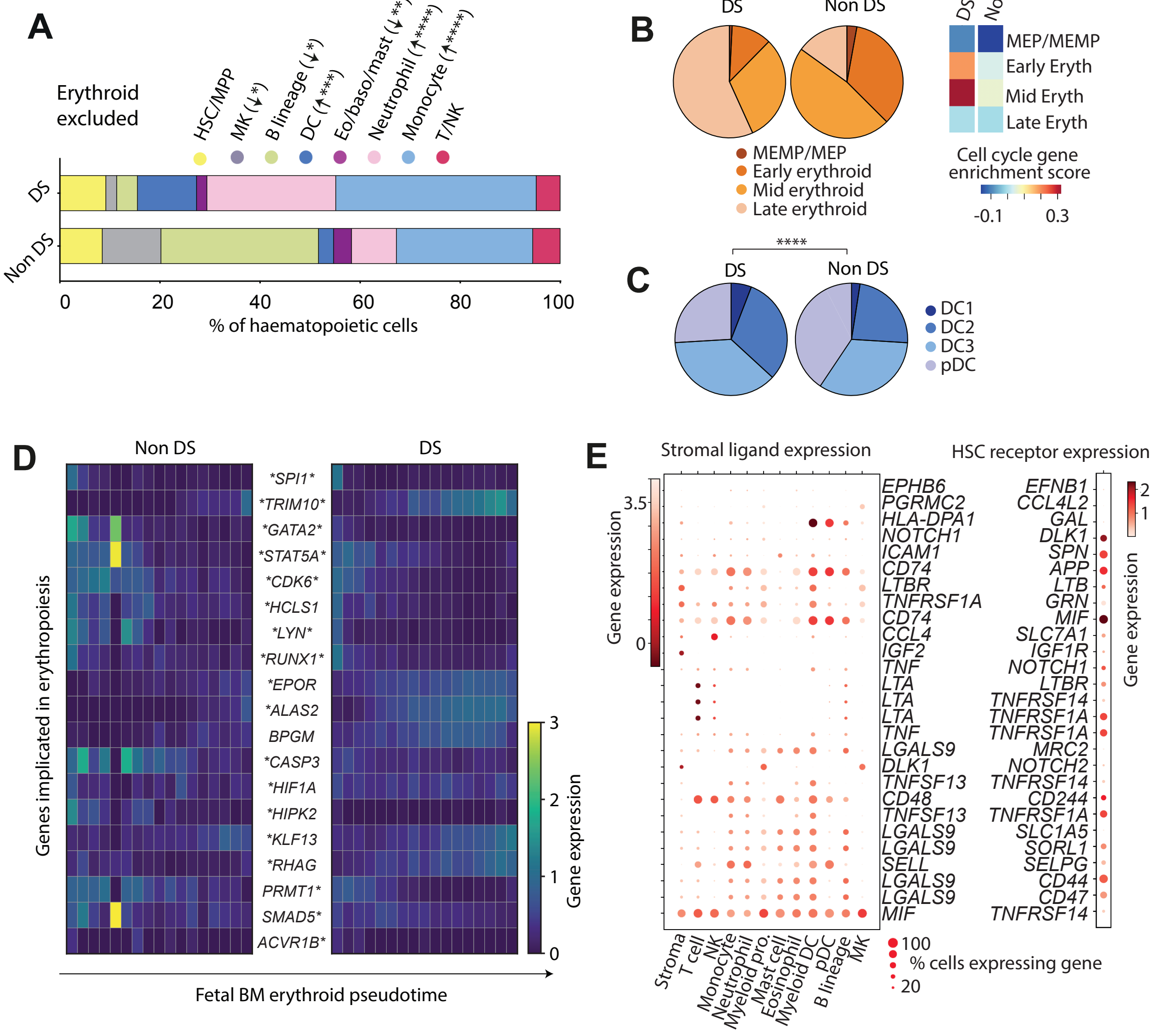

**Supplementary Figure 5: Perturbed haematopoiesis in Down syndrome**

(**A**) Stacked barplot showing percentage of cell types, after excluding the erythroid lineage, in DS (k=2,278) and non DS fetal BM stage 1 (12 - 13 PCW with k = 6,679). Statistical significance of cell frequency difference is shown in parentheses (negative binomial regression with bootstrap correction for sort gates; *$p$ < 0.05, **$p$ < 0.01, ***$p$ < 0.001, ****$p$ < 0.0001).

(**B**) Left panel: Pie chart showing proportions of erythroid cell states in DS vs. non-DS fetal BM. ****$p$ < 0.0001 by Chi-square test. Right panel: heat map showing cell cycle gene enrichment score score in DS and non-DS fetal BM erythroid compartment cells. Relative enrichment is indicated by colour scale.

(**C**) Pie chart showing proportions of DC cell states in DS vs. non-DS fetal BM. ****$p$ < 0.0001 by Chi-square test.

(**D**) Heat maps showing expression of genes implicated in erythropoiesis across Monocle-inferred erythroid differentiation pseudotimes in non-DS and DS fetal BM. Expression values are log-transformed, normalized and scaled. Genes differentially expressed across pseudotime are marked with an asterisk: on the left for non-DS and right for DS.

(**E**) Dot plot showing expression of all ligands significantly overexpressed in DS fetal BM vs. non-DS and the expression of their putative interacting HSC receptor (as inferred by CellPhoneDB). Dotplot constructed as per Fig. 1D legend and methods

### Supplementary Figure 6

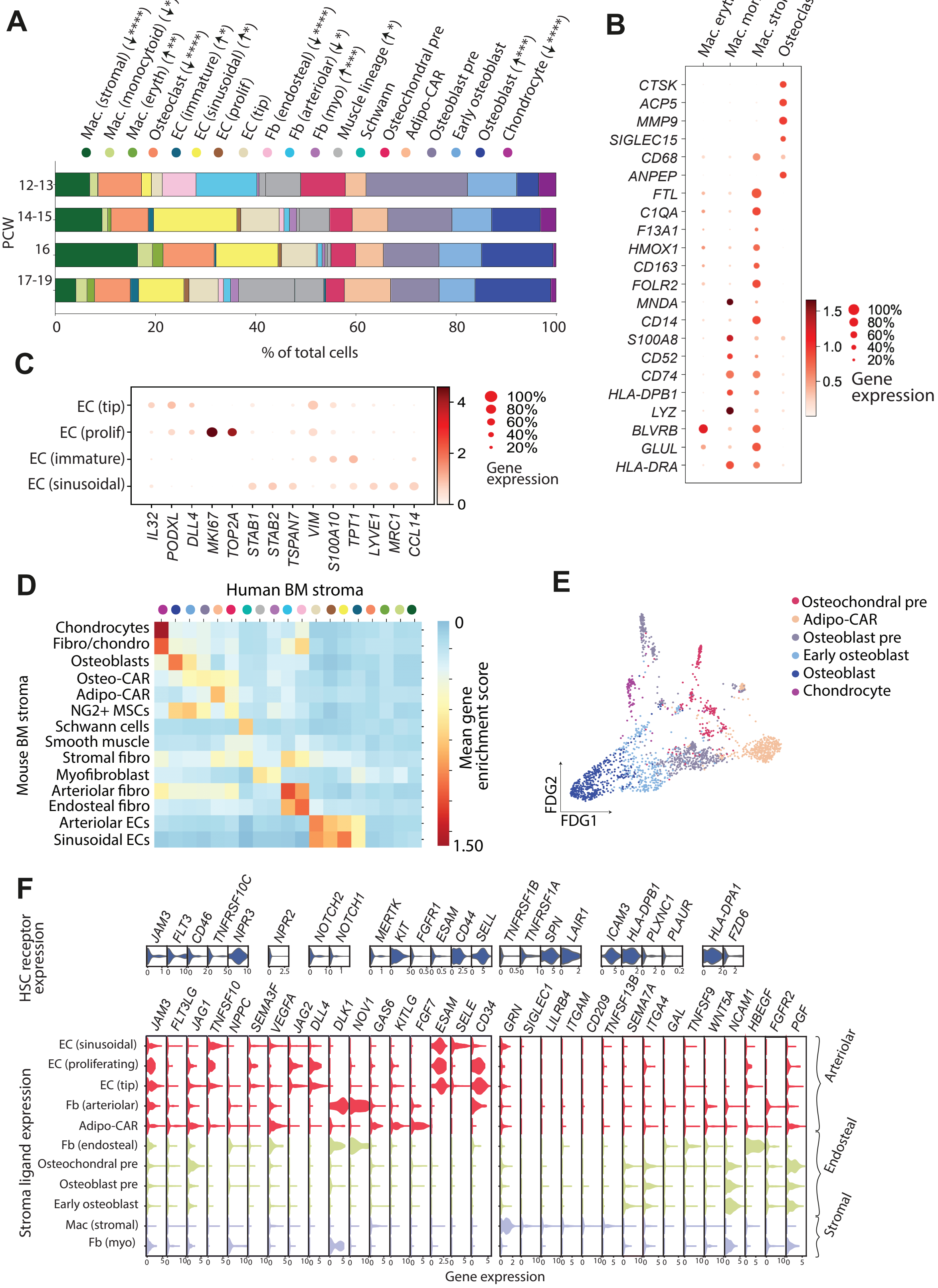

**Supplementary Data Figure 6: Haematopoietic microenvironment in fetal BM**

(**A**) Stacked barplot showing percentage of stromal cell types present in fetal BM across gestational stages. Statistical significance of cell frequency change by gestational stage shown in parentheses (negative binomial regression with bootstrap correction for sort gates; *$p < 0.05$, **$p < 0.01$, ***$p < 0.001$, ****$p < 0.0001$)**.**

(**B**) Identification of distinct osteoclast and macrophage cells states in fetal BM shown by dot plot of cell-state defining marker gene expression. Dotplot constructed as per Fig. 1D legend and methods

(**C**) Identification of 4 distinct ECs in fetal BM states shown by dot plot of cell-state defining marker gene expression.

(**D**) Heat map showing transcriptional similarity between BM stromal cells in post-natal mouse[15] and. and human fetal datasets. Gene enrichment scores are based on gene signatures pooled from top 100 statistically significant DEGs for stromal cell populations in mouse BM[15].

(**E**) FDG visualisation of osteochondral lineage fetal BM cells (k = 1,760).

(**F**) Fetal BM stromal ligands signalling to HSC receptors. Violin plots showing expression of receptor-ligand pairs predicted by CellPhoneDB to have significant interactions in fetal BM scRNA-seq. Log-transformed, normalised and scaled gene expression value is represented on the y-axis.

#### Methods

##### Sample preparation

*Fetal bone marrow tissue acquisition*

Human developmental tissues were obtained from the MRC–Wellcome Trust-funded Human Developmental Biology Resource (HDBR; http://www.hdbr.org) with written consent and approval from the Newcastle and North Tyneside NHS Health Authority Joint Ethics Committee (08/H0906/21+5). HDBR is regulated by the UK Human Tissue Authority (HTA; www.hta.gov.uk) and operates in accordance with the relevant HTA Codes of Practice.

*Fetal developmental stage assignment and chromosomal assessment*

Developmental age was estimated from standardized measurements of foot length and heel-to-knee length[39]. Quantitative Fluorescence PCR of chromosomes X, Y, 13, 15, 16, 18, 21 and 22 was performed on fetal skin or chorionic villi to assign gender and exclude common chromosomal abnormalities. In DS samples, GATA-1s mutation was excluded as previously described[40].

*Dissociation of fetal bone marrow tissue*

Adherent material was removed from fetal femur and bone was cut into small pieces before grinding with a pestle and mortar. Flow buffer (PBS containing 5% (v/v) FBS and 2 mM EDTA) was added to reduce clumping. The suspension was filtered with a 70µm filter then centrifuged for 5 min at 500g. The supernatant was removed before cells were treated with 1x RBC lysis buffer (eBioscience) for 5 min at room temperature and washed once with Flow Buffer before counting.

*Flow cytometry and FACS for scRNA-seq*

Up to 1 million cells were stained with antibody cocktail, incubated for 30 minutes on ice, washed with flow buffer and resuspended at 10 million cells per ml, with DAPI (Sigma-Aldrich) added to a final concentration of 3µM immediately before FACS. FACS was performed on a BD FACSAria Fusion instrument running DIVA v.8 to formulate and execute sort decisions, and data were analysed post-sorting using FlowJo (v.10.6.2, BD Biosciences). For 10x sequencing, cells were sorted into 500µl PBS in pre-chilled FACS tubes coated with FBS (Thermo Scientific). For Smart-seq2, index sorting was used to isolate single cells into 96-well LoBind plates (Eppendorf) containing 10µl lysis buffer (TCL (Qiagen) + 1% (v/v) β-mercaptoethanol) per well. Plates were centrifuged at 300g for 10 seconds and snap-frozen on dry ice for storage at -80° until further processing.

| **10X enrichment** | **Antibody** | **Clone** | **Manufacturer** |
|---|---|---|---|
| | CD45 APC-H7 | 2D1 | BD Bioscience |
| **SS2 FACS** | **Antibody** | **Clone** | **Manufacturer** |
| | CLEC9A PE | 8F9 | Biolegend |
| | CD14 PECF594 | MφP9 | BD |
| | CD66B FITC | REA306 | Miltenyi |
| | HLA-DR PERCPCy5.5 | G46-6 | BD |

| | | | |
|---|---|---|---|
| | CD125 BV421 | A14 | BD |
| | CD123 BV480 | 9F5 | BD |
| | CD34 BV605 | 581 | Biolegend |
| | CD11b BV786 | ICRF44 | BD |
| | CD45 BUV395 | HI30 | BD |
| | CD79A APC | HM47 | Biolegend |
| | CD1c AF700 | L161 | Biolegend |
| | CD52 APCCy7 | 4C8 | BD |
| | CD204 BV711 | U23-56 | BD |
| | CD117 PECy7 | 104D2 | Biolegend |
| **Progenitor FACS** | **Antibody** | **Clone** | **Manufacturer** |
| | CD3 FITC | SK7 | BD Bioscience |
| | CD11c BV421 | B-Ly6 | BD Bioscience |
| | CD14 PE Dazzle | HCD14 | Biolegend |
| | CD16 FITC | NKP15 | BD Bioscience |
| | CD19 FITC | 4G7 | BD Bioscience |
| | CD34 APC-Cy7 | 581 | Biolegend |
| | CD38 PerCP-Cy5.5 | HB-7 | Biolegend |
| | CD45 BUV395 | HI30 | BD Bioscience |
| | CD45RA BV510 | HI100 | BD Bioscience |
| | CD49f PE-Cy7 | GoH3 | eBioscience |
| | CD56 FITC | NCAM16.2 | BD Bioscience |
| | CD90 APC | 5E10 | Biolegend |
| **Culture analysis** | **Antibody** | **Clone** | **Manufacturer** |
| | CD15 BUV395 | HI98 | BD Bioscience |
| | CD11b APC | ICRF44 | Biolegend |
| | CD14 APC-Cy7 | HCD14 | BD Bioscience |
| | CD41 FITC | HIP8 | Biolegend |
| | CD45 V500 | HI30 | BD Bioscience |
| | CD56 PE | NCAM16.2 | BD Bioscience |
| | GYPA BV605 | GA-R2 | BD Bioscience |

*Cytospins*

Cells were sorted into FACS tubes containing chilled PBS. Slides were prepared using a Thermo Cytospin 4 cytocentrifuge and ShandonTM coated slides (Thermo, 5991059), dried at room temperature, then fixed with ice-cold methanol and stained using Giemsa (Sigma-Aldrich), according to manufacturer's instructions. Slides were viewed using a Zeiss AxioImager microscope, images taken of 4 fields from n = 3 samples using the 100X objective, and viewed using Zen (v.2.3) as previously described[4].

*Culture experiments*
Culture experiments were performed on paired fetal BM and FL samples (n=3 , 15-17 PCW), and four additional fetal BM samples (n=4, 14-17 PCW). Cryopreserved single cell suspensions were thawed and sorted into HSC/MPP, LMPP/MLP and $CD34^{+}CD38^{mid}$ fractions as previously described[4], **Supplementary Fig. 4**). Single cells were index-sorted into 96 well plates containing MS5 in log phase growth (DSMZ, passage 6–10), using culture conditions as previously described. Single cell colonies were isolated after 14 days, and prepared for flow cytometry as above. Erythroid colonies were identified as $CD45^{-}GYPA^{+}$ ≥ 30 cells, megakaryocyte colonies as $CD41^{+}$ ≥ 30 cells, myeloid colonies as [($CD45^{+}CD14^{+}$) + ($CD45^{+}CD15^{+}$)] ≥ 30 cells, NK colonies as $CD45^{+}CD56^{+}$ ≥ 30 cells. Two-tailed Fisher's exact tests, performed in Prism (v.8.1.2, GraphPad Software), were applied to the numbers of colonies of each type by stage to determine statistical significance in lineage differentiation potential with development.

*10x scRNA-seq*
Cells sorted for 10x scRNA-seq were counted, then 7,000 cells were loaded onto each channel of a Single Cell Chip before loading onto the 10x Chromium Controller (10x Genomics). Reverse transcription, cDNA amplification and sequencing libraries were generated using either the Single Cell 3' v2 or Single Cell 5' with V(D)J Reagent kits (10x Genomics) as per the manufacturer's protocol. Libraries were sequenced using an Illumina HiSeq 4000 with v.4 SBS chemistry. For the gene expression libraries the following parameters were used: Read 1: 26 cycles, i7 index: 8 cycles, i5 index: 0 cycles, Read 2: 98 cycles. For the V(D)J libraries the following parameters were used: Read 1: 150 cycles, Read 2: 150 cycles. All libraries were sequenced to achieve a minimum of 50,000 reads per cell.

*Plate-based scRNA-seq*
Plates containing lysed single cells were processed using a modified Smart-seq2 protocol[23]. Libraries were generated using the Nextera XT kit (Illumina) with 384 cells per library. Cells were barcoded using Index v.2 sets A, B, C and D (Illumina). Libraries were sequenced using an Illumina NextSeq 550 on High-output mode to achieve a minimum of 1 million reads per cell.

#### Data analysis

*Alignment, quantification and quality control of scRNA-seq datasets*
The fetal BM datasets described in this study (non-DS and DS) underwent pre-processing as detailed below. 10x droplet-based sequencing data was quantified with the CellRanger Single Cell Software Suite (10x Genomics, Inc) and aligned to a GRCh38 human reference genome. Smart-seq2 sequencing data was aligned with STAR (version 2.7.3a) using the STAR index and aligned to the GRCh38 human reference genome. Gene-specific read counts for Smart-seq2 data were calculated using HTSeq-count (version 0.10.0). Cells in scRNA-seq data with fewer than 200 detected genes and genes expressed in fewer than 3 cells were removed from downstream analysis. The methodology for incorporation of external datasets (including: YS, FL, adult BM, CB, blood, thymus, mouse BM) can be found in *Statistics*

*and Reproducibility*, with methods used for any re-annotation described in *Dimensional reduction, visualisation and clustering.*

*Doublet exclusion and transformation of gene expression matrices*
We ran Scrublet (version 0.2.1) on each 10x lane independently, obtaining per-cell scrublet scores. A doublet exclusion threshold of the median plus three times the median absolute deviation scrublet score was applied, as described previously[4]. To alleviate skewness of data and mean-variance relationship, raw gene expression counts were transformed using the *log1p* function in Scanpy (version 1.4.4) in Python (version 3.6.4). To correct for cell-to-cell variation, expression data was then normalised using the *normalize_per_cell* function in Scanpy. Expression values of each gene were then scaled and centred using the *scale* function in Scanpy. Highly variable genes were detected using the *highly_variable_genes* function in Scanpy, with minimum and maximum cut-off values were set as 0.0125 and 3, respectively.

*Dimensional reduction, visualisation and clustering*
Principal components were calculated using the *pca* function in Scanpy. Principal components were adjusted for sequencing type variation (i.e., 3' and 5' sequencing platforms) using the harmony package for batch correction (version 1.0) in R (version 3.6.2). Dependent upon the plateau observed in the elbow curve from the *pca_variance_ratio* function in Scanpy, an informative number of PCs were selected for downstream analysis. The *neighbours* function in Scanpy was used to calculate the neighbourhood graph. Uniform manifold approximation and projection (UMAP) embedding was calculated using the *umap* function in Scanpy. The neighbourhood graph was then clustered using the *leiden* function in Scanpy. Gene expression dot plots were visualised using the Scanpy package in Python with dot colour indicating the mean logged, normalised and scaled expression values and dot size representing the proportion of cells annotated to each category that express the given gene.

*Annotation of clusters*
Cluster cell identity was assigned through analysis of DEGs and their alignment with marker genes identified through literature search. DEGs were calculated in Scanpy using the *rank genes_groups* function, which performed a Wilcoxon rank sum test restricted to genes expressed in at least 25% of cells in either of the two populations compared, and with a log (natural) fold change cut-off of 0.25. All p-values were adjusted for multiple testing using the Benjamini-Hochberg method. Following initial broad rounds of annotations, clusters of broad similarity (e.g., lymphoid cells) were subset for further rounds of feature selection, visualisation, clustering and annotation as described above. Clusters whose gene signatures indicated additional diversity were further investigated in an iterative manner, and those unique signatures were selected for downstream analysis, leading to greater resolution and confidence in cell type annotation.

*Functional profiling of differentially expressed genes*
Cluster cell identity was reinforced by passing DEGs through functional profiling using clusterProfiler (version 3.14.3). Analysis of a given cluster's enriched pathways, Gene Ontology (GO) terms and KEGG (Kyoto Encyclopaedia of Genes and Genomes) terms further enforced confidence in annotations based on DEGs and known cell type markers. Analysis of a given cluster's enriched pathways, GO terms and KEGG terms were performed using the compareCluster function in the clusterProfiler package in R.

clusterProfiler dot plots were output with dot colour indicating statistical significance of the enrichment (q value) and dot size representing the fraction of genes annotated to each term.

*Gene enrichment scores*

To conduct gene enrichment scores against a reference published blood dataset [23], the top 100 DEGs (log2 fold change) of blood DC and monocyte cell types were input into *sc.tl.score_genes* function in Scanpy. Gene enrichment value for the blood reference cell type was then calculated as the average expression of the top DEG from the reference dataset, minus the average expression of another reference set of genes (randomly sampled from each binned expression value). Gene enrichment scores were visualised using a heatmap in the seaborn (v.0.9.0) Python package.

Cell cycle gene enrichment scores were calculated through use of a curated list of genes implicated in the human cell cycle [41] as the input for *scanpy.tl.score_genes_cell* function in Scanpy. The G2/M and S phase score for each cell thus represented low to high enrichment for a particular phase's genes. In order to serve as a proxy for a 'proliferative phase' score, the mean of the G2/M and S phase scores were calculated and plotted in UMAP space. Cells with G2/M/S cell cycle score greater than the mean were assigned as 'cycling' cells, else assigned as 'not cycling'. Gene enrichment scores calculated using *sc.tl.score_genes* function in Scanpy were used to ascertain i) apoptotic gene enrichment through use of genes implicated in the KEGG apoptotic pathway (GSEA: M8492) and ) NK cytotoxic gene enrichment through use of genes implicated in the KEGG NK cytotoxicity pathway (GSEA: M5669).

*Calculating differences in cell type proportions across gestational stages and organ*

GraphPad Prism v8.1.0 was used for plotting and statistical comparison. Statistically significant differences in cell type proportions across tissue were conducted using a one-way ANOVA with Tukey's multiple comparison tests. Significance was noted on corresponding scatter plots using asterisks, where scatterplots display proportion per biological replicate. Cell-type proportions per sample were obtained by adjusting observed proportions by $CD45^+$/$CD45^-$ sort gate. For cell type proportion statistical analysis across gestational stage and tissue, proportions were modelled as a quasibinomial distribution. For both analyses the condition (gestational stage or tissue) was provided as a covariate for the proportion of the cell type being assessed. The quasibinomial model was fit using glm from the MASS R package. The p-value for the significance of the change in proportion between conditions was assessed using a likelihood-ratio test, computed using the anova function. Significant changes in cell type proportion were highlighted on bar plots using asterisks. The Spearman's rho test was used to assess monotonic trends of flux between analogous cell states of different developmental stages. Increasing or decreasing trends were denoted with "up" and "down" arrows respectively.

*VDJ analysis*

Using the pyVDJ Python package (v0.1.2), lanes of BCR-enriched and TCR-enriched 10x data were integrated with their corresponding 10x GEX lane data in Scanpy. Filtered CellRanger output files were then imported into the Scanpy workflow to investigate productivity of chains, presence of heavy and light chains and clonal assignment. VDJ metadata by cell type was then exported from Scanpy and plotted in GraphPad Prism.

*Trajectory inference using Monocle3*
Gene expression matrices for cell-types of interest (filtered by highly variable genes, as defined in Scanpy) were loaded into the Monocle workflow as CellDataSet objects using Monocle3 version 0.2.1 in R. Gene expression values were then normalised by log and size factor to address depth differences using *preprocess_cds* function. For known lineages, cells were clustered with a resolution parameter of 1e-07 in order that one partition was returned (to ensure pseudotime with incorporated all cells). Cells were then ordered along pseudotime and with root state provided using the *order_cells* function. DEGs across pseudotime were calculated using a Moran's I statistical test (*graph_test* function) and DEGs grouped into 'modules' by their Moran's-derived correlation across pseudotime using the *find_gene_modules* function. Dynamically expressed genes across a given pseudotime were then plotted as heatmaps, with normalised logged and scaled gene expression values. Paired heat maps across DS/non-DS and across tissue were the product of combined processing (log, normalising, scaling) of Gene expression counts and plotting gene expression over independently derived pseudotime trajectories.

*Prediction of cell-cell communication using CellPhoneDB*
To assign putative cell-cell interactions within our fetal BM scRNA-seq dataset, we used CellPhoneDB v2.1.2 to identify significant receptor ligand interactions between stromal cells and progenitor cell types. Log, normalised and scaled expression values for cell types of interest were exported from Scanpy along with their respective cell type metadata. Using database version 2.0.0, CellPhoneDB was run using the statistical method, with *p*-value cut-off of 0.05 for significant receptor ligand pairs and a result precision of 3dp. To summarize the data for visualization, cells were aggregated into groups: arteriolar (EC-sinusoidal, EC-proliferating, EC-tip, EC-arteriolar), endosteal (Fb-endosteal, Osteochondral precursor, Early osteoblast) and stromal (Mac-stromal, Fb-fibroblast). Total cell numbers of interest were included in analysis unless down-sampling stated in *Statistics and reproducibility.*

*Inference of transcription factors and their gene regulatory networks using PySCENIC*
The PySCENIC package (v0.9.19) and pipeline was used to identify transcription factors and their target genes in the combined no DS and DSs datasets. The ranking database (hg38__refseq-r80__500bp_up_and_100bp_down_tss.mc9nr.feather), motif annotation database (motifs-v9-nr.hgnc-m0.001-o0.0.tbl) and list of transcription factors (lambert2018.txt) were downloaded from the Aert's laboratory github page. An adjacency matrix of transcription factors and their targets was generated and pruned using the Aert's group suggested parameters. Comparisons of TF activity between no DS and DS were made using t-tests of the AUCell output of predicted transcription factor activity in each cell for each cluster. The regulons generated were used to predict which genes controlled by each transcription factor in downstream analysis.

*Network analysis and clustered pathway annotation*
The FindConservedMarkers function in Seurat V3.1 (bonferroni corrected FDR adjusted P values < 0.05) and a Benjamini-Hochberg-corrected Wilcoxon rank-sum test (log fold change > 0.25 and *p*-values < 0.05) were used to identify conserved and differentially expressed genes between analogous cell states in each dataset. Genes were submitted for over-representation analysis (ORA) using the G-profiler2 package

in R to query two databases simultaneously (Reactome, Gene Ontology (GO) Biological Process). We derived statistical significance ($q < 0.05$) for each gene set enrichment and performed Markov clustering (MCL) using the MCL package in R to derive network neighbourhoods based on shared genes between the gene sets. The gene set clusters were annotated using the AutoAnnotate Cytoscape package. Clusters were ranked by the mean enrichment score of all gene sets within each cluster and manually curated based on biological significance. We used Cytoscape to visualise clusters of enriched gene sets. Transcription factor regulation of each markov cluster was identified by hypergeometric modelling of genes in the clustered genesets to a pre-compiled TF association matrix acquired from the Enrichr database of ENCODE and ChEA consensus TF associations using the Hypergeometric package in R, over-represented TFs were then ranked and filtered by *p*-value (<0.05). To characterise inflammatory response pathways associated with the stroma in DS fetal BM, we derived DEGs between analogous stromal compartments in DS and non-DS and submitted the genes for network analysis as outlined prior. Fold changes for genes associated with the top five Markov clusters ranked by cluster size were visualised with the ggplot2 (3.3.2) package in R.

*TNF response gene annotation*
We derived DEGs between analogous cell state compartments in DS and non-DS fetal BM, statistically significant DEGs (*p*-value < 0.05) were compared for intersect against TNF response associated genes acquired from the GO biological process database (GO:0034612). Intersecting TNF response genes in DEGs were ranked by log fold change between cell states of DS and non-DS.

**Statistics and reproducibility**
*Fetal liver, fetal yolk sac and thymus scRNA-seq data*
For all re-analysis of FL and YS *scRNA-seq* data (n=113,063 and 10,071), n=14 and 3 biologically independent samples were used and total population of annotated cell types were shown in figures unless otherwise stated. Original cell numbers can be found in the original publication[4].

For **Fig. 2B**, proportions of myeloid cell states arising from BM haematopoiesis and their counterparts in other tissues were compared. Macrophages were not included in this analysis. The YS cell state originally assigned as DC progenitor in [4] was renamed as macrophage, after further exploration and re-annotation, and therefore not included here. Further YS myeloid states - GMP, promonocyte and MOP were identified on re-clustering the Lymphoid progenitor, MEMP, Myeloid progenitor and YS progenitor/MPP. For **Fig. 2B**, the original FL Monocyte precursor and Neutrophil-myeloid progenitor were subclustered to resolve heterogeneity and revealed further myeloid states including MOP, monocyte, promonocyte and promyelocyte cells. FL myeloid nomenclature was updated so that monocyte precursor became promonocyte and monocyte became $CD14^+$ monocyte. Due to the presence of unique pDC precursor in FL, pDC and pDC precursor were merged into one grouping cross-tissue for purposes of bar plot. For **Fig. 4D**, the original fetal YS Lymphoid progenitor, MEMP, Myeloid progenitor and YS progenitor/MPP in [4] were sub clustered to identify HSC, MEMP, GMP, CMP, ELP and MEP. For **Fig. 4D**, the FL HSC_MPP, MEMP and Pre pro B cell in [4] were sub clustered to further identify HSC, MEMP, GMP ELP, MPP, MEP, eo/baso/mast precursor, myeloid DC progenitor and pDC progenitor. In **Fig. 6B** FL endothelial populations were re-clustered to annotate sinusoidal endothelium. For all analysis of thymus 10x data (n=259,265) n=24 biologically independent samples were used and total population of annotated cell types were shown in figures, with exception of down-sampling in **Fig. 3G**, where 1,000 thymus DN cells were used for analysis. Cell numbers can be found in the original publication[3,4].

*Fetal bone marrow scRNA-seq data*
For analysis of combined fetal BM 5'and 3' data (n=104,562), n=9 biologically independent samples were used. Total population of annotated cell types were shown in figures unless otherwise noted.

In **Fig. 3B**, only B cells with corresponding VDJ data (5,052/28,613) were shown in the bar plot. In **Supplementary Data Fig. 3F**, only single positive T cells with corresponding VDJ data (138/560) were shown in barplots. In **Supplementary Data Figs. 4A, B**, the CMP, ELP, GMP, MEP, MOP, early MK, early erythroid, eo/baso/mast precursor, pre pro B progenitor and promyelocyte were downsampled to a population of 70 cells each. For **Fig. 4D**, the fetal BM progenitor compartment comprised those cells grouped into the HSC_MPP broad annotations. In **Supplementary Data Fig. 5D**, non-DS fetal BM MEP and MEMP were merged and all non-DS fetal BM cells were down-sampled to include only 12-13 PCW samples (matched age with DS fetal BM). For CellPhoneDB analysis in **Fig. 5E**, DS and non-DS fetal BM cells were combined for input to CellPhoneDB, with non-DS fetal bone marrow down-sampled to match cell numbers in fetal BM DS.

The following refined cell number annotations were displayed in each of the figures: CD4 T cell - 327, CD8 T cell - 171, CD14 monocyte - 8787, CD56 bright NK - 450, CMP- 425, DC1 - 50, DC2 - 598, DC3 - 705, DC precursor - 201, EI macrophage - 92, ELP - 1358, GMP - 1285, HSC - 92, ILC precursor - 67, LMPP - 34, MEMP - 16, MEP - 269, MK - 1036, MOP - 3990, MPP myeloid - 92, NK T cell - 111, NK progenitor - 26, Treg - 62, adipo-CAR - 359, arteriolar fibroblast - 84, basophil - 139, chondrocyte - 81, early MK - 1665, early erythroid - 7534, early osteoblast - 291, endosteal fibroblast - 54, eo/baso/mast precursor - 175, eosinophil - 325, erythroid macrophage - 92, immature B cell - 1998, immature EC - 69, late erythroid - 4649, mast cell - 648, mature NK - 136, mid erythroid - 14408, monocytoid macrophage - 296, muscle - 219, muscle stem cell - 255, myelocyte - 3854, myeloid DC progenitor - 31, myofibroblast - 78, naive B cell - 1423, neutrophil - 4516, osteoblast - 375, osteoblast precursor - 463, osteochondral precursor - 191, osteoclast - 1378, pDC - 713, pDC progenitor - 23, pre B progenitor - 14234, pre pro B progenitor - 5428, proliferating EC - 26, promonocyte - 7676, promyelocyte - 2386, schwann cells - 9, sinusoidal EC - 550, stromal macrophage -1493, tDC - 193, tip EC - 363, pro B progenitor - 5530.

When cell types were grouped into broad lineages (e.g., **Figs. 1B, C**), cell numbers were as follows: HSC_MPP - 3800, erythroid - 26591, MK - 2701, B_lineage - 28613, DC - 2460, eo/baso/mast - 1112, neutrophil - 10756, monocyte - 20453, T_NK - 1350, stroma - 6726. Other broad groupings are detailed in supplementary tables.

*Fetal bone marrow Smart-seq2 data*
For analysis of fetal bone marrow Smart-seq2 validation data (n=486), n=2 biologically independent samples were used and the following cell numbers were shown: $CD34^+$ = 32, B cell = 52, DC1 = 34, DC2 = 15, monocyte = 32, PMN = 65, basophil = 20, eosinophil = 54, mast cell = 47, myelocyte = 61, pDC = 30, promyelocyte = 44.

*Fetal bone marrow Down syndrome scRNA-seq data*
For all analysis of DS fetal BM 5' data (n=8,662), n=4 biologically independent samples and total cell populations were used, unless stated otherwise.

The following refined cell number annotations were displayed in each of the figures: CAR - 4, CD8 T cell - 55, CD14 monocyte - 252, CD56 bright NK - 36, CMP - 37, DC1 - 16, DC2 - 94, DC3 - 98, HSC - 45, ILC precursor -13, MEMP - 42, MK - 25, MOP - 350, MSC -18, Treg - 8, chondrocyte - 4, early B cell - 23, early MK - 7, early erythroid - 766, endothelium - 37, eo/baso/mast precursor - 40, eosinophil - 41, late erythroid - 3341, macrophage - 64, mast cell - 27, mature B cell - 20, mature NK - 69, mid erythroid - 2082, myelocyte - 218, neutrophil - 245, osteoblast -11, osteoclast - 17, pDC - 14, pre B cell - 59, promonocyte - 321, pre pDC - 40, promyelocyte - 72, transitional NK cell -11.

When cell types were grouped into broad lineages (e.g., **Fig. 5A, B**), cell numbers were as follows: HSC/MPP = 231, Erythroid = 6189, MK = 32, B lineage = 102, DC = 262, Neutrophil = 820, Eo/baso/mast = 68, Monocyte = 571, TNK = 192, Stroma = 155. Other broad groupings are detailed in supplementary tables.

*Adult bone marrow scRNA-seq data*
For all analysis of adult BM scRNA-seq (n=142,026) data, n=4 biologically independent samples were used, unless stated otherwise. Lanes from four donors (BM1, BM2, BM5, BM6) were downloaded from the Immune Cell Atlas public repository (https://data.humancellatlas.org/). For **Fig. 4D**, the adult BM progenitor compartment comprised those cells grouped into the HSC_MPP broad annotations.

The following refined cell number annotations were displayed in each of the figures: CD14 monocyte - 3670, CD16 monocyte - 1938, CD56 bright NK - 1228, CLP - 882, CMP - 288, DC1 - 135, DC2 - 481, DC3 - 550, DC precursor - 462, HSC - 497, LMPP - 80, MEMP - 785, MK - 577, MOP - 1440, MPP - 365, Treg - 6327, early MK - 136, early erythroid - 5441, erythroid macrophage - 77, immature B cell - 2728, late erythroid -1150, mature CD8 T cell - 15725, mature NK - 6074, memory B cell - 4106, memory CD4 T cell - 22197, mid erythroid - 2192, monocyte-DC - 515, myelocyte - 6675, myeloid DC progenitor - 110, naive B cell - 19265, naive CD4 T cell - 5873, naive CD8 T cell - 8965, neutrophil - 2482, pDC - 1134, pDC progenitor - 63, plasma cell - 2074, pre B cell - 971, pro B progenitor - 1390, promonocyte - 7448, promyelocyte - 2197, stroma - 161, tDC - 75, transitional B cell - 2151, transitional NK - 946.

When cell types were grouped into broad lineages (e.g., **Supplementary Data Fig. 1E**), cell numbers were as follows: HSC/MPP = 3,007, Erythroid = 8,783, MK = 713, B lineage = 32,685, DC = 3,415, Neutrophil = 11,354, Monocyte = 14,496, TNK = 67,335, Stroma = 238.

*Cord blood scRNA-seq data*
For all analysis of CB 10x (n=148,442) data, n=4 biologically independent samples were used, unless stated otherwise. Lanes from four donors (CB1, CB2, CB5, CB6 respectively) were downloaded from the Immune Cell Atlas public repository (https://data.humancellatlas.org/). For **Fig. 4D**, the CB progenitor compartment comprised those cells grouped into the HSC_MPP broad annotations.

The following refined cell number annotations were displayed in each of the figures: CD8 T cell - 16345, CD14 monocyte - 13324, CD16 monocyte - 888, CD56 bright NK - 4066, CMP - 272, DC1 - 67, DC2 -155, DC precursor - 169, GMP - 203, HSC - 194, ILC precursor -1519, MEMP - 338, MK - 1262, early MK - 496, early erythroid - 532, late erythroid - 878, mature NK - 7860, mid erythroid - 2627, myelocyte - 3726, naive B cell - 19516, naive CD4 T cell - 69338, neutrophil - 3458, pDC - 242, preDC - 269, promonocyte - 607, tDC - 91.

When cell types were grouped into broad lineages (e.g., **Supplementary Data Fig. 1F**), cell numbers were as follows: HSC/MPP - 1007, erythroid - 4037, MK - 1758, B cells - 19516, DC - 993, neutrophil - 7184, monocyte - 14819, T/NK - 99128.

*Blood monocyte and DC scRNA-seq data*
Monocyte-DC blood scRNA-seq (SS2) data were downloaded from [23]. The available RPKM counts for 1140 monocytes (n=768) and DCs (n=372) were log-transformed and scaled in preparation for DEG analysis conducted as described below.

*Differentially expressed gene statistics*
Differential gene expression analysis referenced in text and shown in violin plots were run using the Wilcoxon rank-sum statistical test with Benjamini-Hochberg procedure for multiple testing correction. *p*-values are shown in the relevant supplementary tables.

*Statistics from barplots*
For cell type proportion analysis across gestational stages and different tissues, proportions were modelled as a quasibinomial distribution. *p*-values for the significance of change in proportion between conditions were assessed and values provided in figure legends. For all barplot statistics, asterisks were used to indicate significant changes in proportion, with *, **, *** and **** representing *p*-values of <0.05, <0.01, <0.001 and 0.0001 respectively.

*Statistics from colony experiments*
Statistical analysis of culture wells producing colonies between fetal BM and FL HSC/MPPs by Mann Whitney test yielded p=0.006 (***, n=7 experiments). Comparison between fetal BM and FL committed progenitors by the same method yielded *p*=0.006 (***, n=1536 wells). Comparison of number of colony types per well for paired progenitor types was performed by binomial test, comparing 1 colony type with >1 colony type. *p*-values were 0.0012 for fetal BM and FL HSC/MPPs (**, n=164) and 0.3566 for fetal BM and FL committed progenitors (ns, n=152). Comparison of number of myeloid-only colonies for paired fetal BM and FL HSC/MPPs was performed by binomial test, comparing ‘myeloid only’ with ‘myeloid+other’. *p*-values were 0.0002 (***, n=79).
